# Marine heatwaves threaten cryptic coral diversity and erode associations amongst coevolving partners

**DOI:** 10.1101/2023.01.07.522953

**Authors:** Samuel Starko, James Fifer, Danielle C. Claar, Sarah W. Davies, Ross Cunning, Andrew C. Baker, Julia K. Baum

## Abstract

Climate change-amplified heatwaves are known to drive extensive mortality in marine foundation species. However, a paucity of longitudinal genomic datasets has impeded understanding of how these rapid selection events alter species’ genetic structure. Impacts of these events may be exacerbated in species with obligate symbioses, where the genetics of multiple co-evolving species may be affected. Here, we tracked the symbiotic associations and fate of reef-building corals for six years through a prolonged heatwave. Coral genetics strongly predicted survival of the common coral *Porites* through the event, with strong differential survival (15 to 64%) apparent across morphologically identical -but genetically distinct- lineages. The event also disrupted strong associations between coral lineages and their symbiotic partners, homogenizing symbiotic assemblages across lineages and reducing the specificity of coral-algal symbioses. These results highlight that marine heatwaves threaten cryptic genetic diversity of foundation species and have the potential to decouple tight relationships between co-evolving host-symbiont pairs.

## Introduction

Extreme climatic events, including heatwaves, wildfires, floods, and droughts, are now major agents of change, posing serious threats to biodiversity and natural ecosystems globally (*1–3*). Such events are driving factors behind many species range contractions, extirpations, and invasions (*4–8*), and may also be influencing evolutionary processes through selection imposed by rapid environmental change (*9–11*). Recent studies have demonstrated that selection through extreme events can be directional, favoring individual genotypes carrying adaptive traits (*10, 12– 14*). While this selection may help species survive future events of a similar nature, it also threatens to reduce overall genetic diversity, which may limit species’ capacity to respond to new selection pressures, such as those imposed by extreme climatic disturbances of different durations or intensities (*15*), or to unexpected stressors and pathogens (*16*).

In the ocean, some of the most profound impacts of climate change are experienced during marine heatwaves (*5, 17*) – pulse heat stress events in which water temperatures are abnormally high for unusual lengths of time (*18, 19*). While it is well documented that heatwaves can threaten marine biodiversity by driving conspicuous species losses (*4, 5, 20*), selection imposed during heatwaves may also drive cryptic losses of genetic diversity within taxa (*9, 10, 12*), a phenomenon that is predicted to be an outcome of these events but has only rarely been demonstrated in marine systems due to the lack of baseline genetic data (*10, 12*). Losses of genetic diversity could have long-term evolutionary consequences for taxa by limiting their scope for future adaptation (*10, 12, 21*). However, remaining populations may also have increased heat tolerance due to directional selection, possibly improving the fitness of future populations in a warming ocean (*10*). Our ability to anticipate these evolutionary consequences will depend on understanding the extent to which marine heatwaves drive differential mortality among genotypes (i.e., natural selection), altering the genetic structure of marine taxa (*10*). To date, the few studies that have been done in this area have demonstrated that marine heatwaves can alter the relative abundance of different genotypes (*12*), but linking these patterns to fitness components, which requires tracking survival and/or reproductive output of individuals, remains a challenge (but see *22*).

Tropical coral reefs are now considered as the most vulnerable coastal marine ecosystem in the face of climate change (*23*), with marine heatwaves their primary threat. Heatwaves disrupt the critical relationship between reef-building corals and their obligate endosymbionts (family Symbiodiniaceae), causing them to bleach (*17, 24, 25*) and making them vulnerable to starvation and disease (*26*). Intense or prolonged marine heatwaves can cause mass bleaching and widespread coral mortality, with profound ecological and socioeconomic impacts (*17*). This is especially true given that these ecosystems are among the most biologically diverse and economically valuable in the ocean (*27*). Corals may primarily adapt to climate change through either shifts in host allele frequencies through adaptation (*28*) or shifts in their microbial symbiont communities (*29–31*), which are heritable to varying degrees (*32*). Yet, while a handful of studies have tracked the stability of symbioses through marine heatwaves and shown differential bleaching by algal symbiont (*11, 33, 34*), only one has directly assessed the impacts of these events on the population genetics of coral taxa in natural systems (*22*). Moreover, despite growing awareness that cryptic coral genotypes can harbour unique assemblages of symbionts, which could be the primary determinants of their climate change vulnerability (or resilience) (e.g., *33, 35*), no study to date has simultaneously tested for shifts in both host population genetics and associated symbiont assemblages through an extreme heatwave event. Thus, the extent to which heatwaves drive differential mortality or alter patterns of symbiont specificity across co-occurring coral genotypes, potentially threatening rare or heat-sensitive lineages of either symbiotic partner, remains largely unclear (*11, 36*).

As in a wide range of taxa, molecular investigations of reef-building corals over the past two or more decades have drastically reshaped our understanding of their evolution and diversity (e.g., *25, 37, 38*). Similar to macroalgae and other invertebrates (e.g., *39, 40*), many morphologically-defined coral species actually represent cryptic species complexes consisting of multiple morphologically similar, or indistinguishable, lineages that are partially or completely reproductively isolated from one another (e.g., *38, 41–43*). Although coral cryptic lineage complexes are common, the number of studies testing for differences in heat tolerance between lineages is limited (*33, 44*) and, with one recent exception (*22*), have only assessed variation in bleaching tolerance, rather than survival through natural heatwaves (e.g., *33*), despite the fact that these two processes can be decoupled (*34*). Quantifying the strength of selection on corals and their obligate symbionts through marine heatwaves is essential to understanding and predicting the influence of future heatwaves on the genetic diversity and adaptive potential of threatened coral reefs.

Between 2014 and 2017, a series of heatwaves unfolded across much of the world’s tropical reefs (*17, 45*). This period, considered chronologically as the 3^rd^ global coral bleaching event on record, was unprecedented in terms of the severity, duration, and geographic spread (*45*). This event led to mass coral bleaching and mortality across many coral reefs in the Pacific and Indian Oceans, including extensive damage to the Great Barrier Reef (*17, 34, 46, 47*). Species-level assessments of coral mortality have demonstrated that there were winners and losers in the face of this widespread bleaching, with survival varying substantially across coral taxa (*46, 48*). However, it is not known whether selection imposed by mass mortality during this global bleaching event impacted the genetic composition of coral species or populations, nor how underlying local anthropogenic stressors – a feature of virtually all coral reefs – might modulate impacts on this critical facet of diversity.

Here, we directly assessed the extent to which marine heatwaves drive differential mortality across coral genotypes and alter the specificity of host-symbiont pairings. We focused on one of the most widespread, ecologically significant, and well-studied coral genera, *Porites*, and tracked the fate and algal symbiont composition of individual *Porites* colonies (massive growth-form; field identified as *Porites lobata*) in the central equatorial Pacific Ocean, through the 3^rd^ global coral bleaching event. Within this region, the coral atoll Kiritimati experienced some of the highest levels of accumulated heat-stress ever documented on a coral reef, rivaled only by the nearby Jarvis Island during this same time period (*48*). This heatwave lasted ten months, imposing ∼31.6 degree heating weeks (°C-weeks) on Kiritimati’s coral reefs (*34*). Despite this, massive *Porites* had relatively high survivorship (∼80% at some sites), with highly variable bleaching severity and survival among colonies and sites (*48*). We leveraged this extreme climatic event as a natural experiment to directly test whether coral bleaching susceptibility or survivorship could be predicted by the genetic structure of the affected colonies and/or their associated algal symbionts, and further assessed changes in the relative abundance of host and symbiont genotypes by comparing a larger sampling of colonies from before, during and after the heatwave. Recent molecular studies have determined that the genus *Porites* comprises at least eight clades, some characterized by complex genetic structure (*41, 49*), possibly reflecting cryptic or pseudo-cryptic lineages within each clade. Our study focused on one of these clades (Clade V from ref *49*; also known as the *Porites lobata/lutea clade*)) facilitating a deeper look into the functional differences between finer-scale cryptic lineages than has previously been achieved in this group. Our objectives were to examine if 1) cryptic coral lineages were present, and if they differed in their ability to survive a marine heatwave; 2) if this differential survival was modulated by underlying exposure to chronic local human disturbance; 3) if cryptic coral lineages were associated with specific symbionts; and 4) whether the specificity of these symbiotic partnerships was impacted by mass bleaching and mortality during the heatwave.

## Results

### *Sympatric cryptic lineages of* Porites

We identified three genetic lineages of massive *Porites* (hereafter referred to as PKir-1, PKir-2, and PKir-3) that were found sympatrically across the reefs of Kiritimati prior to the 2015-2016 El Niño-driven heatwave (Fig. 1). Ordination (based on >12,000 SNPs from 2b-RAD) revealed three distinct genomic clusters with no intermediate genotypes, which was further supported by ADMIXTURE analyses showing the lowest CV error for k = 3 where every sample was assigned to a lineage with >85% probability (Fig. 1a,b). Global F_ST_ values between lineages were also high, suggesting relatively high levels of differentiation across cryptic lineages (Table S1). PKir-1 and PKir-2 (Global F_ST_ = 0.263) were found to be more genetically similar to each other than either was to PKir-3 (Global F_ST_ = 0.361 and 0.326, respectively; Fig S1). However, historical gene flow was found between all lineages (Fig S2), suggesting that, although these lineages appear reproductively isolated in the present day, they have likely experienced introgression in the past.

**Fig 1.**
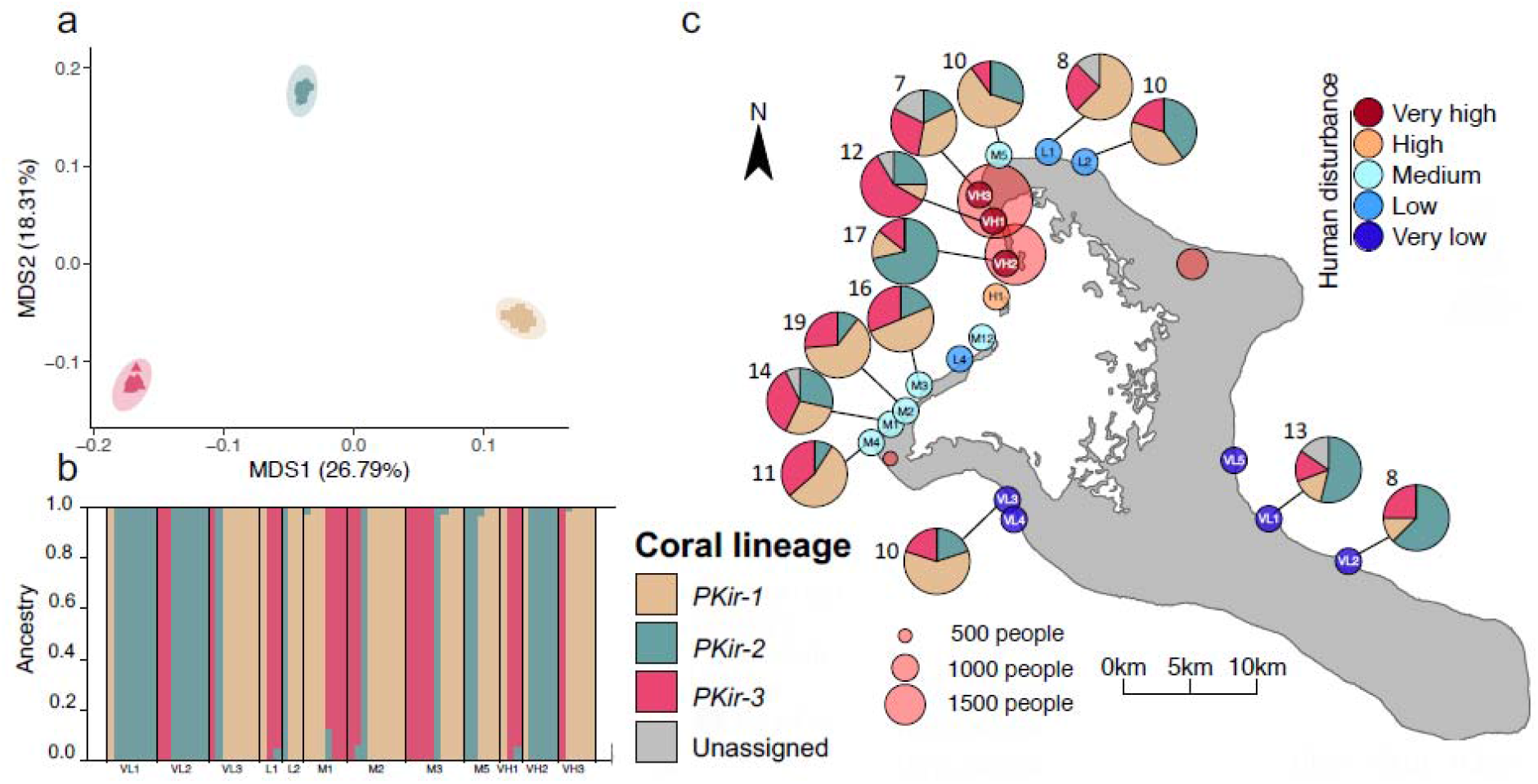
Cryptic lineages of massive *Porites* across forereef sites on Kiritimati. (a) Principal Coordinate Analysis (PCoA) of 2b-RAD data (using 1-Pearson correlation matrices through ANGSD) showing three population clusters. (b) Results of ADMIXTURE analysis showing the assignment of colonies to one of three lineages, arranged by collection site. (c) A map with pie charts showing the relative abundance of each lineage at each site before the heatwave. Numbers indicate the number of colonies sampled and sequenced with either 2b-RAD or ITS2 metabarcoding. Circles indicate sites colored by level of human disturbance and scaled by human population size.

Demographic analyses infer some limited gene flow between lineages with asymmetrical introgression across lineages and regions of the genome. The best-fit model supported the hypothesis of heterogeneous gene flow across the genome, with a small proportion of the genome experiencing particularly high gene flow (Fig S2). Moreover, we inferred higher gene flow from PKir-3 to PKir-1 and PKir-2 compared to the reverse direction (Fig S2). Effective population sizes (N_e_) were similar across all three lineages, and all showed contraction in recent millennia (Fig S3).

Leveraging host sequences in the ITS2 metabarcoding dataset, we were able to expand lineage assignment beyond those colonies that were sequenced with 2b-RAD (n = 67). As expected, all *Porites* ITS2 sequences belonged to one of the 8 previously described *Porites* lineages (clade V or the *Porites lobata/lutea* clade (*49*); Fig S4). However, examining colonies that were sequenced using both ITS2 and 2b-RAD (n = 64 with successful host sequences), we found that ITS2 sequences were consistently dissimilar across cryptic *Porites* lineages. The set of sequences found in each cryptic lineage was paraphyletic relative to the sequences of other lineages (Figs S4, S5, Table S2), likely reflecting the recent reticulate divergence of these cryptic lineages and confirming that this clade consists of a cryptic complex rather than a single, highly plastic species (see discussion in *41*). Moreover, several colonies sampled with both ITS2 and 2b-RAD (n = 12) were heterozygous at the ITS2 locus, with two dominant host sequence variants identified from each colony. In total, 23 ITS2 sequence variants were present across the 64 colonies sequenced with both metabarcoding and 2b-RAD. Two sequence variants were found in both PKir-1 and PKir-2, making them uninformative for lineage assignment. However, these sequence variants were relatively rare across the dataset (ASV9: 19/305 - 6% of colonies; ASV31: 4/305 – 1% of colonies). Several uncommon ITS2 sequence variants (n = 10) were not found in any samples assigned using 2b-RAD, preventing lineage assignment in these cases. In total, we were able to assign 92% (281/305) of colonies included in this study using either 2b-RAD or ITS2 metabarcoding data. Using all samples collected prior to the heatwave for which lineage assignment was possible (n = 149), we found that, although there was a relationship between the relative abundance of each lineage and region of the atoll (Bayes Factor [BF] = 36.10), there was no relationship with local human disturbance (BF = 0.84).

### Survivorship through a heatwave varies by cryptic lineage and human disturbance

Tracking individual colonies through nine time-points that span the 2015-2016 heatwave, we found strong evidence of differential survival across lineages, but only at sites without very high levels of human disturbance (Fig. 2). Mortality of tagged colonies began during the heatwave (first observed in May 2016) and continued for several months following the heatwave with no mortality observed after 2017. While mortality was generally much higher at sites with increased human disturbance, survivorship up to 2017 depended on the interaction between human disturbance and lineage, with PKir-3 having only ∼15% survival across all disturbance levels, and PKir-1 and PKir-2 having ∼70-90% survival at minimally disturbed sites, but only ∼5% survival at the most disturbed sites (Logistic regression; lineage*disturbance: P = 0.007). There was also a significant effect of cryptic lineage identity on bleaching score, such that PKir-3 tended to have the highest level of bleaching at both time-points during the heatwave (2015: Deviance = 9.329, p = 0.010; 2016: Deviance = 9.412, p = 0.009; Fig S6).

**Fig 2.**
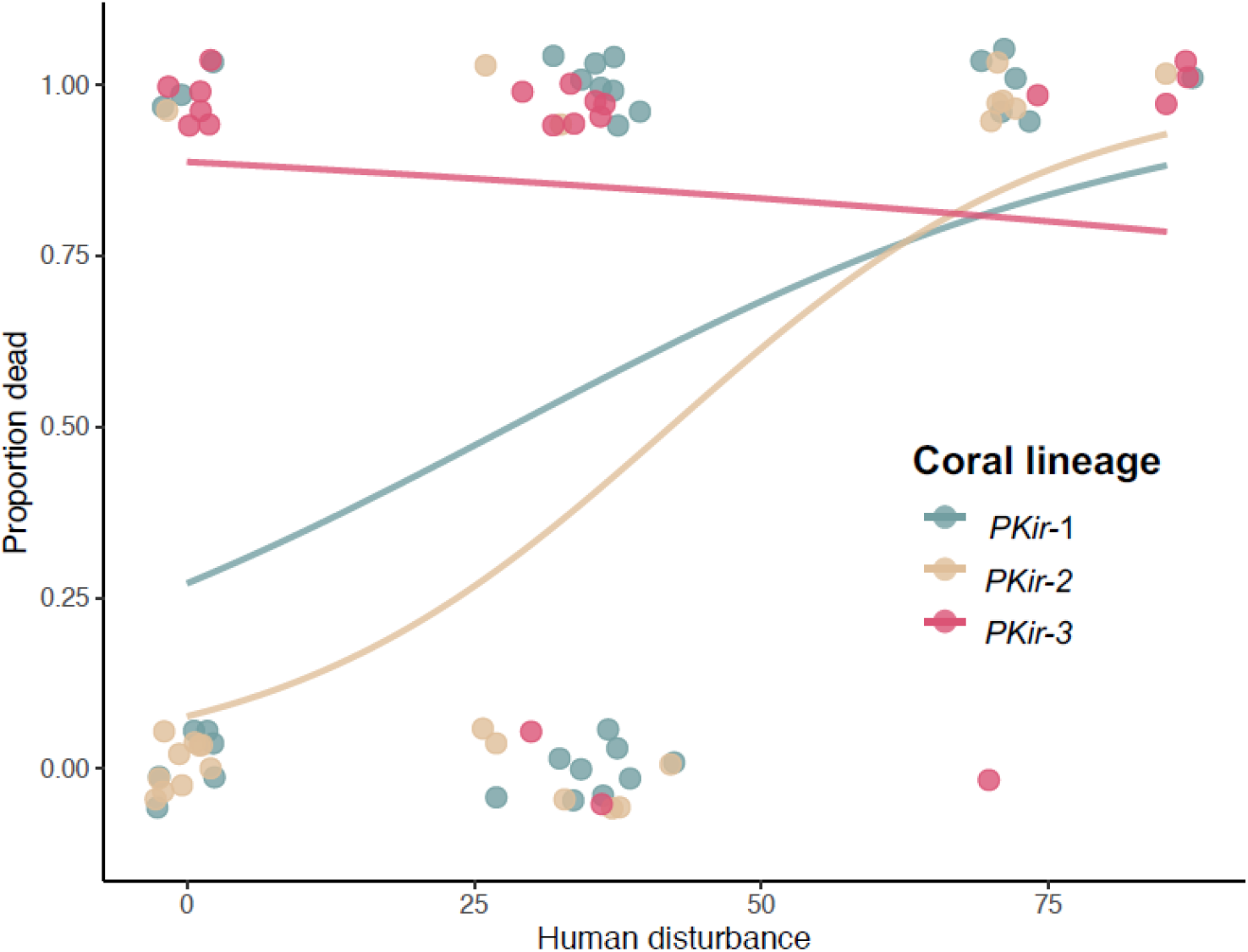
Survivorship by coral cryptic lineage and chronic human disturbance. Each point represents a colony that either survived (0) or died (1). The proportion of colonies that died at each value is estimated by the logistic regression line. Note that human disturbance is a relative metric based on fishing pressure and distance to Kiritimati’s villages (see ref., *72*). Note that data points are jittered for visualization.

We tested for local genomic differentiation across the three *Porites* lineages and identified genes near outlier loci. While we found several genes near outlier loci when comparing lineage pairs (PKir-1 vs PKir-2: n = 47; PKir-1 vs. PKir-3: n = 63; PKir-2 vs. PKir-3: n = 42; Supp File 1), the only gene near an outlier locus when comparing both PKir-1 and PKir-2 to PKir-3 matched the ETS-related transcription factor Elf-2 (∼57% similarity).

### Disruption of lineage-specific symbioses

We found strong associations between coral lineage and symbiont assemblage composition prior to the marine heatwave, but these associations were disrupted following the event. Across all colonies sampled before the heatwave, there was a strong relationship between coral lineage and recovered *Cladocopium* sequence variants from the C15 clade (PERMANOVA: F = 175.41, R^2^ = 0.73, P < 0.001). Specifically, sequences from the C15 clade formed at least two clusters (Fig 3A), with variants in one cluster associating almost exclusively with PKir-3 colonies (all but one case, ∼2.5%, although two PKir-3 colonies, 5%, also had symbiont sequences from the other cluster). For colonies sampled after the heatwave, this association between symbiont sequence variants and coral lineage was disrupted, with sequences from all lineages forming a single cluster (Fig. 3B). After the heatwave, only a single colony of unknown lineage (due to lack of host sequence reads) was found to still possess sequence variants common in PKir-3 prior to the heatwave.

**Fig 3.**
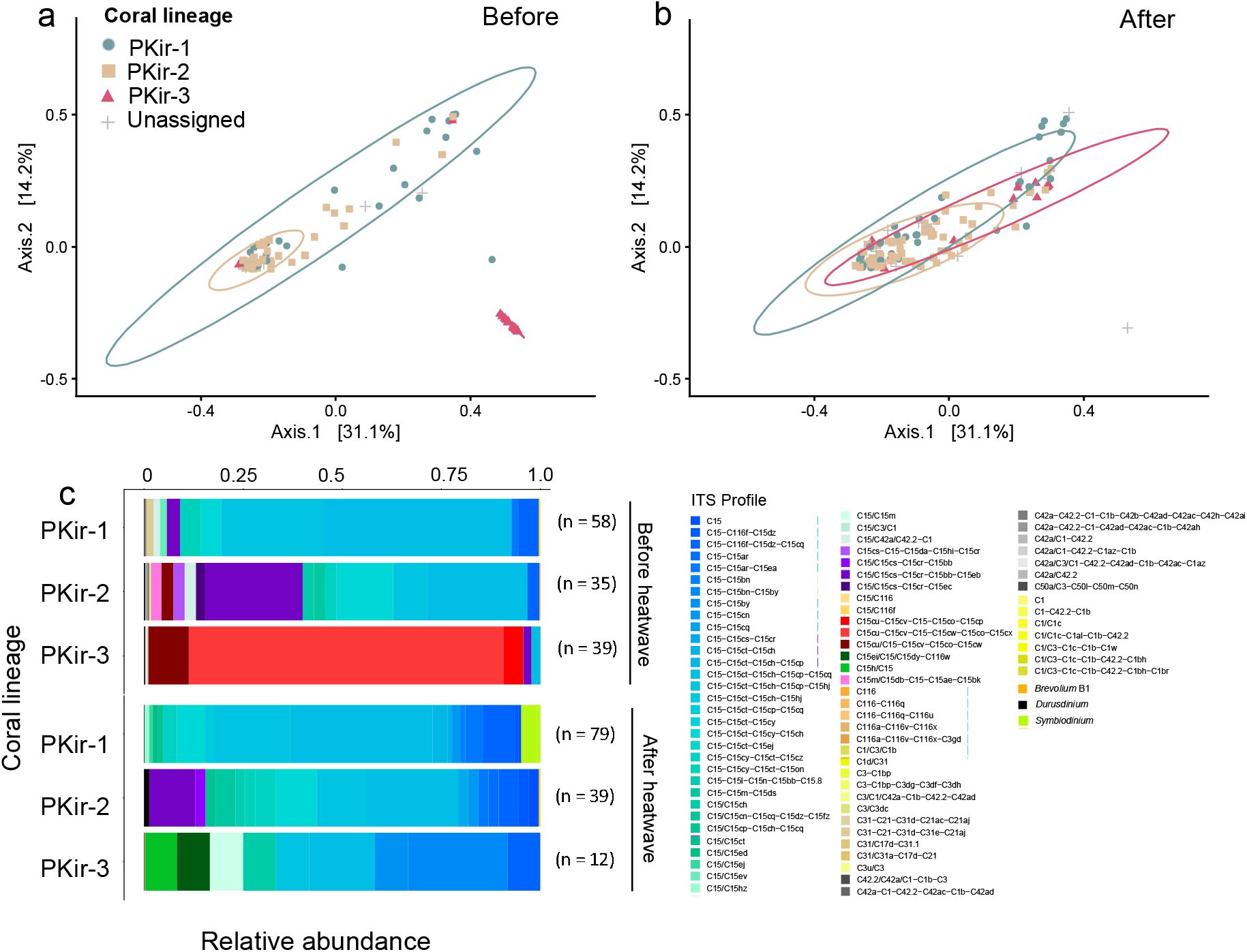
Impact of the marine heatwave on lineage-specific symbioses. Shown are the results of PCoA (based on Unifrac distance) of *Cladocopium* C15 sequence variants for all colonies sampled (a) before and (b) after the heatwave. Ellipses at the 95% level are shown for each assigned coral lineage. Also shown in (c) are the relative abundances of each ITS2 profile across all colonies of each lineage before and after the heatwave. Sample sizes indicate the number of colonies. Only samples with > 500 sequence reads were included. Note that the ellipse for PKir-3 in panel a largely overlaps the points and is therefore challenging to visualize.

These patterns in algal symbiont communities were also captured by ITS2 profiles, a means of characterizing symbiont types that attempts to identify putative Symbiodineaceae taxa (*50*) (Fig S7). Nearly all corals sampled at any time point (629/653 samples; 96%) were characterized by a single *Cladocopium* profile each from the C15 clade. A small percentage (∼2%) of colonies had mixed assemblages that included a single profile each from the C15 radiation and one or two additional profiles from other Symbiodiniaceae lineages (e.g., *Cladocopium* C116, *Durusdinium* D1, D4). In total, we identified 45 profiles from the C15 radiation (113 profiles from all Symbiodineaceae lineages) across all colonies successfully sequenced. No corals had more than one profile from the C15 clade and only ∼1 % of samples lacked a profile from the C15 clade altogether (mostly from March 2016; see below). Unlike PKir-1 and PKir-2, which initially associated with 9 and 13 profiles, respectively, PKir-3 had highly specific symbiotic associations prior to the heatwave with 95% of colonies (37/39) associated with one of just three symbiont profiles within the C15 radiation that was nearly absent from the other lineages (PKir-1: 0%, PKir-2: 3%) (Fig S8). These tight associations broke down during the heatwave such that these profiles were completely absent from PKir-3 colonies sampled after the heatwave (n = 12; Fig 4c).

**Fig 4.**
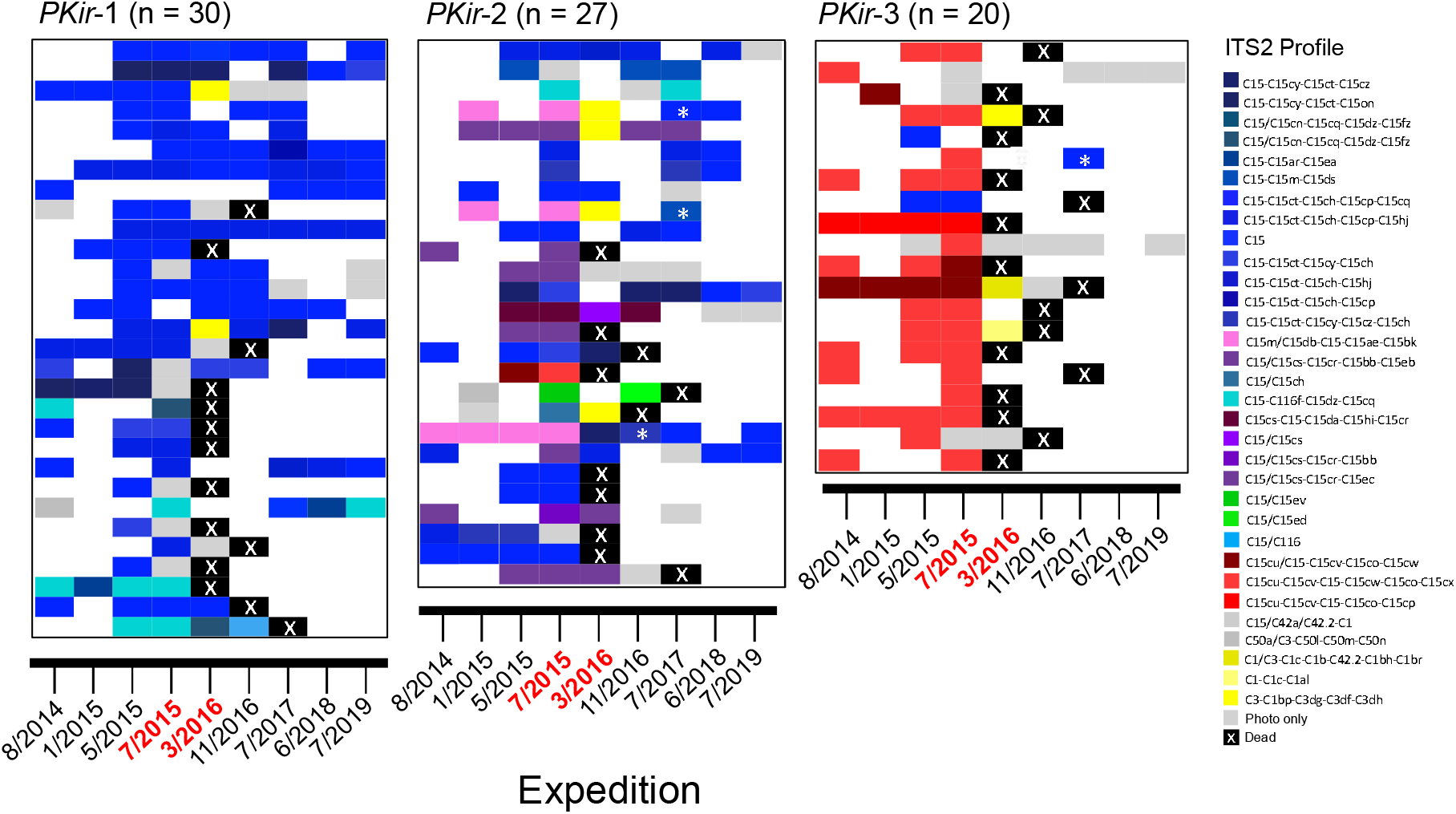
Temporal stability of Symbiodiniaceae associated with tracked colonies of each cryptic *Porites* lineage. Each row represents an individual colony, with colour at each time point indicating the dominant symbiont profile. Colonies within each lineage are arranged in order of human disturbance (lowest on top, highest on bottom). Note that colonies that survived to 2017 were considered alive for survivorship analyses. Colonies that were considered to have switched or shuffled profiles from the C15 clade (i.e., between ITS2 profiles with different dominant DIVs; n = 4) are indicated with an asterisk. Expeditions during the heatwave are shown in red text.

Although most of the colonies that were tracked for two or more time-points (∼64%; 100/157) had the same profile from the C15 clade in every case, approximately one third of colonies did host variable symbiont profiles over time (e.g., see Fig. 4). Most notably, a few colonies sampled both before and after the heatwave appeared to recover from bleaching with a new profile from the C15 lineage. For example, one of the three surviving colonies of PKir-3 switched from a “C15cu” profile (i.e., dominated by the C15cu sequence variant) to a “C15” profile (dominated by the C15 sequence variant). The other two PKir-3 colonies that survived were not successfully sequenced with ITS2. Although, it is possible that these colonies still possess “C15cu” profiles, this would still represent a shift in the relative abundance of “C15” profiles in PKir-3 from ∼5% to >80% across the lineage at large. Similarly, three PKir-2 colonies switched from “C15m” (which were previously only found in that *Porites* lineage) to “C15” profiles, and symbiont assemblages across PKir-3 colonies, in general, became more homogenous, with decreases in the relative abundance of “C15m” and “C15/C15cs” profiles in favour of “C15” profiles (Fig 4, Fig S7). Some additional colonies also switched between very similar profiles (with the same dominant sequence but having additional minor sequence variants; 39/157). The overall similarity of these latter profiles suggest they may represent closely-related members of the same symbiont population that may have even been assigned as different profiles due to sequencing artifacts (e.g., missing a rare sequence variant).

We also identified several cases where bleached colonies contained high relative levels of three profiles from *Cladocopium* C3 or C1 symbiont lineage, suggesting these symbionts may be residual or opportunistic in these *Porites* hosts. These three symbiont profiles (1 - C1/C3-C1c-C1b-C42.2-C1bh-C1br; 2-C1-C1c-C1al; 3-C3-C1bp-C3dg-C3-df-C3-dh) only appeared in bleached colonies during the March 2016 expedition, which was late in the heatwave (Fig. 4). Three of these colonies survived and were resampled during a later expedition; all three had reverted to hosting symbionts from the C15 clade following recovery from bleaching. Other symbiont types (e.g., C116 or *Durusdinium*) were generally rare and found inconsistently across samples, suggesting that these are minor or opportunistic constituents of the *Porites* holobiont; they were not further examined here. However, we note that these rare and/or transient profiles may be of functional importance to the *Porites* holobiont, possibly even facilitating the shuffling or switching of dominant profiles from the C15 lineage.

## Discussion

By coupling host genomic sequencing and Symbiodiniaceae metabarcoding with longitudinal coral colony tracking, we have demonstrated that cryptic lineages of *Porites* coral experienced strong differential mortality during a tropical marine heatwave of unprecedented duration. We identified three distinct lineages of massive *Porites* and tracked colonies of each lineage through ten months of intense heat-stress, demonstrating much higher mortality in one lineage, PKir-3, than in the other two. Human disturbance modulated this effect of host lineage, with a strong relationship between disturbance and mortality in the PKir-1 and PKir-2 lineages, and no difference in lineage-specific mortality observed at sites exposed to *very high* disturbance levels, where high mortality was observed for all corals regardless of their lineage. This differential selection resulted in a substantial change in the relative abundances of these cryptic lineages, with the relative abundance of PKir-3 decreasing by 70% across the atoll following the heatwave (30% to 9% relative abundance of tagged corals of known lineage).

Cryptic *Porites* lineages also differed in their associated symbiotic ITS2 profiles but only prior to the 2015-2016 bleaching event. *Porites* corals generally transmit their symbionts vertically such that symbiont genotypes are heritable across generations of coral (*51*). This could lead to strong patterns of phylosymbiosis and/or cophylogeny (*52–54*), where closely related corals share similar algal symbiont communities, a pattern clearly reflected across host lineages in our dataset.

Indeed, differences in symbiotic assemblages between cryptic lineages have now been documented across multiple coral genera (*33, 35, 42*). Under strong selection from the heatwave, however, this pattern of co-occurrence between coral host lineage and algal symbiont sequence variants was disrupted in our study. Following the heatwave, we only found a single sample containing “C15cu” symbionts (from a colony of unknown lineage), even though these symbionts were found in 95% of PKir-3 colonies before the heatwave (and almost 25% percent of colonies overall). All confirmed post-heatwave samples of PKir-3 (though not all sampled prior to the heatwave) were instead associated with “C15” symbionts rather than “C15cu” profiles. Among the tracked colonies, four colonies (three from PKir-2 and one from PKir-3) that bleached and recovered post-heatwave did so with different profiles from the C15 lineage than they hosted prior to bleaching (Fig 4). The erosion of this pattern of phylosymbiosis across coral lineages is likely driven by some amount of symbiont switching or shuffling that occurred during recovery from bleaching. However, given that these profiles were present in ∼5% of PKir-3 colonies before the heatwave, differential mortality across colonies with differing profiles from the C15 clade may have also played an important role in driving these observed patterns.

Although we cannot definitively tease apart the impacts of symbiont identity from other genomic factors, the breakdown of patterns of host-symbiont associations and the observed switching of symbionts in some colonies are most parsimoniously explained by functional differences between the ITS2 profiles within the C15 phylotype. This hypothesis is further supported by the fact that all sampled colonies that were ‘healthy’ late in the heatwave (i.e., May 2016) were associated with “C15” profiles including the two healthy colonies from PKir-3 (which more typically associated with “C15cu” and did not remain healthy). Although functional differences between Symbiodineaceae genera are well documented (i.e., *Cladocopium* vs. *Durusdinium* (*55, 29, 31, 34*)), it has remained unclear until recently whether closely related sequence variants (i.e., C15 variants) can express functional variation that is meaningful in the face of heat stress. However, recent work showed that closely related variants of C15 were associated with bleaching variation between *Porites cylindrica* and *Porites rus* (*56*) – two clearly defined (i.e., both morphologically and genetically) species. Moreover, variants of C3 in the Persian Gulf have rapidly evolved increased thermal tolerance relative to close relatives from nearby areas (*57*). Our study offers supporting evidence to the hypothesis that closely related algal symbionts can vary substantially in function by demonstrating how intense warming can result in the near complete loss of a previously prominent symbiont genotype while increasing the relative abundance of its close relatives.

Massive *Porites* were initially assumed by some authors to only inherit their symbionts vertically and have fixed symbiont dominance (*58*). However, multiple studies have now shown that a single massive *Porites* colony can harbour mixed *Cladocopium* and *Durusdinium* communities (*59, 60*) as well as different profiles from the C15 lineage (*43*), suggesting the ability for either “shuffling” of dominant symbionts (*29*) or horizontal transmission of new Symbiodiniaceae (*61*). Indeed, *Porites* can harbour different dominant ITS2 profiles across environmental gradients, suggesting that symbiotic variation has ecological implications for host colonies (*62, 63*). Our data confirm the hypothesis that *Porites* can shuffle or switch symbionts by demonstrating their ability to shift between profiles from the C15 clade following extreme bleaching. The ability to shuffle or switch symbionts may be adaptive by allowing corals to avoid evolutionary “dead-ends”, whereby a vertically transmitted symbiont is fixed across the host population, but may be maladaptive under future warming (*24, 55*). Bleaching and shuffling or switching symbionts, however, came at a great cost to the population size of PKir-3 with a mortality rate in that lineage exceeding 80%. Lab experiments on *Montipora*, which also transmits its symbionts maternally, have shown that changes in symbiont assemblages acquired in one generation can be transferred to the next (*32*), providing an avenue for intergenerational plasticity in coral holobiont function (*28*). This suggests that the loss of variation in symbiont identity across colonies may have long-term consequences for the range of symbiont-host pairs found on Kiritimati which, in turn, may reduce the functional diversity and adaptability of corals facing future warming. In contrast, however, the remaining colonies of PKir-3 may now be better adapted to heatwave events of similar nature, if their newly dominant symbionts increase their thermal tolerance in the face of future events (*55*). Overall, our results demonstrate how a single extreme event can decouple potentially tight co-evolutionary relationships between symbiotic partners.

In 2016, late in the heatwave, several of the bleached colonies of all lineages were associated with “C1” and “C3”-dominated ITS2 profiles that were only ever present during that expedition (∼10 months into the heat stress; Fig 4). In all cases, these colonies were severely bleached when sampled and surviving colonies recovered “C15” sequence variants during later time points (see Fig 4). Thus, due to the transient nature of C1 and C3 profiles, we interpret these associations as opportunistic Symbiodiniaceae infections. However, it is also possible that these are very rare, residual profiles that remained following bleaching and were not detected prior due to the much higher relative abundance of other profiles during non-bleached timepoints. While it remains unclear whether these opportunists offered any benefit to the corals, it is possible that they helped colonies maintain some basic nutritional requirements during the period between initial bleaching and subsequent recovery of symbionts from the C15 lineage. A similar pattern was observed during bleaching of *Pocillopora* spp. in the eastern Pacific, where bleached colonies were temporarily colonized by an opportunistic *Breviolum* population (*11*). The functional and ecological importance of these short-lived symbioses remain unclear but offer an interesting avenue for future research.

Differences in survival across lineages likely reflect differences in the timing of bleaching. Late in the heatwave, most corals had experienced some bleaching and many were severely bleached.

However, PKir-3 had the highest proportion of bleached colonies early in the heatwave and had far fewer ‘healthy’ colonies later in the heatwave compared to the other two lineages. This suggests that the increased mortality in PKir-3 was a result of these colonies spending a longer time bleached, perhaps the consequence of less thermally tolerant symbionts. We did, however, also identify one gene that was an outlier between both PKir-3-PKir2 and PKir3-PKir1 genomic comparisons, ETS-related transcription factor Elf-2, which may have possible links to coral immunity (*64*). Thus, it is also possible that genetic differences between these cryptic lineages influenced the probability of survival, for example, by increasing bleaching propensity or susceptibility to disease following bleaching. However, this hypothesis remains highly speculative.

Although all three lineages of *Porites* were sympatric across Kiritimati, the extent to which they fully overlap across the seascape remains unclear. Past work on cryptic lineages has demonstrated that they often occur in slightly different habitats even if they do overlap in geographic distribution (e.g., *33, 43*). Although currently unclear, asymmetrical gene flow that we inferred across lineages could be the result of differences in habitat. For example, if PKir-3 is found across a larger range of habitats than the other two lineages, then this could help to explain why gene flow was reduced into PKir-3. Given the functional differences in thermal tolerance through a major heat-stress event, it is possible that these lineages occupy different depth ranges, for example, but co-occur in the moderate forereef environment (as observed in *Pocillopora* spp. In Mo’orea (*35*)). Understanding the distribution of cryptic coral lineages across different environments will be important for elucidating the processes driving and reinforcing differentiation across these lineages and better predicting future bleaching events (*38*).

Cryptic lineages are being rapidly discovered across a broad range of taxa (*39*, e.g., *65*) but their functional importance is unclear, particularly when they are distinguished by fine-scale genetic differences. Theory would predict functional differences between cryptic lineages if they have diverged as a result of ecological speciation (*66–68*). However, there is also substantial evidence that close relatives tend to be ecologically similar when comparing to a broader pool of taxa (*69*). Thus, it remains unclear whether climate change has the potential to impose directional selection on these cryptic lineages or whether closely related lineages can instead be expected to respond similarly.

Here, we demonstrate strong differential mortality among cryptic coral lineages during a prolonged marine heatwave, providing direct evidence that heatwaves have the potential to threaten cryptic genetic diversity, even among one of the most common and stress tolerant coral genera. Cryptic lineages had specific symbiont associations that recombined during the heatwave, highlighting a likely mechanism behind differential survival of lineages. Moreover, mortality was strongly predicted by human disturbance in two of the three cryptic lineages, illustrating that anthropogenic drivers can mediate the strength of selection during extreme events. High mortality in PKir-3 decreased its overall population size, increasing the probability that the lineage goes extinct in the near future. However, changes in the symbiont associations of that lineage may facilitate adaptation to future heatwaves, with unknown functional trade-offs (*55*). While a population-wide shift in associated symbionts may be a form of adaptation, increasing colony-level thermal tolerance in the face of future events (*24*), the loss of this specific host-symbiont pairing demonstrates how heatwaves may be eroding biotic interactions in addition to threatening diversity. Nonetheless, our study demonstrates that strong marine heatwaves may drive biodiversity loss at finer scales than have generally been appreciated to date. Overall, these finding underscore the need to better understand genetic diversity within our current conceptions of species. Moreover, they illustrate how climate change may threaten the persistence of undiscovered diversity, causing Centinelan extinctions – losses of taxa that are never described by science and therefore unrecorded (*70*). Moreover, this undescribed diversity is likely to explain meaningful variation in coral bleaching and mortality which has remained challenging to predict.

## Materials and Methods

### Study location and design

Kiritimati (Christmas Island), Republic of Kiribati, is located in the central equatorial Pacific Ocean (01°52′N 157°24′W), at the center of the Niño 3.4 region (a delineation used to quantify El Niño presence and strength). Kiritimati is the world’s largest atoll by landmass (388 km^2^; 150 km in perimeter), and all eighteen surveyed reefs surrounding the atoll are sloping, fringing reefs with no back reef or significant reef crest formations. Kiritimati has a strong spatial gradient of human disturbance, with the majority of the human population restricted to two villages on the northwest side of the atoll. Human uses, including waste-water runoff, subsistence fishing, and a large pier, are densely concentrated in this area, while other parts of the atoll experience substantially less human disturbance. The intensity of chronic local human disturbance at each site has previously been quantified, using two spatial data sources: 1) human population densities and 2) fishing pressure (*34, 48*). First, as a proxy for immediate point-source inputs from villages into the marine environment such as pollution and sewage runoff, a geographic buffer (in ArcGIS) was generated to determine human population size within 2 km of each site. Nearly all people live in villages, and village location was mapped based on published field surveys. Population size for each village was extracted from the 2015 Population and Housing Census from the Kiribati National Statistics Office (*71*). Secondly, to account for the more diffuse effects of subsistence fishing on the reef ecosystem, a kernel density function with ten steps was generated based on mapped fishing intensity from household interviews conducted by Watson et al. (*72*). Each metric was weighted equally, and from this combined metric sites were grouped into five distinct disturbance categories, termed *very low, low, medium high*, and *very high*. When treating human disturbance as a continuous variable, we square root-transformed the combined metric to account for skew in the data (as in *34*).

### Coral tagging and sampling

We tagged and sampled colonies of massive *Porites* along 60 m transects, laid along the 10-12 m isobath, at each of the 18 different fore reef sites around Kiritimati (Fig. 1C), during expeditions before (August 2014, January/February 2015, April/May 2015), during (July 2015, March 2016), and after (November 2016, July 2017, June 2018, July 2019) the 2015-2016 El Niño. All colonies were identified in the field as *Porites lobata*. Twelve of the sites were sampled both before and after the heatwave; one site was sampled before but could not be accessed after, and five of the sites were sampled only after the heatwave. In total, 305 massive *Porites* colonies were included in this study from at least one timepoint on the basis that symbiont and/or host lineage was obtained from sequence data (detailed below). At each visit, each coral colony was photographed and a tissue sample was taken, except in the few cases following the heatwave when the live tissue remaining on the colony was too small to sample.

Of the total set of colonies included in this study (n = 305), 157 colonies were initially tagged and sampled before the heatwave (n= 6 - 20 at 13 sites). However, not all sites could be visited during each expedition, and some site surveys were only partially completed during some expeditions due to unfavourable weather conditions or other logistical constraints. Among these 157 colonies, we were able to track the survivorship of 79 through the mortality event (3 – 10 per site) on the basis that tagged colonies could be relocated at various timepoints spanning the heatwave. All but two of the tracked colonies (n = 77) were assigned to a cryptic *Porites* lineage using either 2b-RAD sequencing or host ITS2 metabarcoding data. Several additional colonies were sampled only during (n = 6), after (n = 100), or during and after (n = 42) the mortality event. Some colonies were sampled but sequences could not be obtained due to sample quality and/or failed benchwork.

Mortality from bleaching began some time following the July 2015 expedition and continued until at least late 2016 but ceased following the July 2017 expedition (Fig 4). Thus, we considered a colony to have died if its mortality occurred between March 2016 and July 2017 and we considered colonies to have survived if they were found alive in 2017 or later. None of the 79 colonies tracked through the heatwave died later than 2017 but some were not tracked beyond that timepoint. Because symbionts remained stable and no mortality had yet occurred at the early time-point in the heatwave, we considered all 2015 surveys to have occurred before the mortality event. These colonies were not included in survivorship analyses but provide additional insight into the relative abundance of different coral lineages and symbiont sequence variants across the various expeditions. Overall, sample sizes of each analysis vary depending on the number of colonies for which necessary information (e.g., host lineage, symbiont sequence variant, survivorship) were available (Table S3-S5).

### Assessing bleaching of tagged colonies

Bleaching was assessed visually from photographs of each colony, using a categorical score based on the percentage of the colony that was visually bleached. We considered colonies with less than 10% bleaching to be “healthy”, those with 11-50% bleaching to have experienced “partial bleaching”, and colonies with >50% bleaching were considered to be “severely bleached”.

### DNA extraction

We performed DNA extraction using one of two methods: 1) a guanidinium-based extraction protocol optimized for Symbiodiniaceae DNA (*30, 73*) and 2) a second protocol using the DNeasy Blood and Tissue kit performed with modifications to optimize for coral genomic DNA extraction (*74*). Following the first protocol, the DNA pellet was washed with 70% ethanol three times rather than once and, if necessary, the final product cleaned using Zymo Genomic DNA Clean and ConcentratorTM-25 (Catalog Nos. D4064 & D4065) following the standard protocol (http://www.zymoresearch.com/downloads/dl/file/id/638/d4064i.pdf). For ITS2 metabarcoding, the guanidinium-based extraction was used. However, for 2b-RAD preparations, coral genomic extractions were used unless there was no remaining tissue left from that sample, in which case the guanidium-based extraction was used.

### High-throughput sequencing

We used amplicon sequencing to characterize the algal symbiont communities associated with each coral colony. We chose the ITS2 amplicon for high-throughput sequencing because it is currently the standard region used for identification and quantification of Symbiodiniaceae taxa (*50*). Library preparation for Illumina MiSeq ITS2 amplicon sequencing was performed following the Illumina 16S Metagenomic Sequencing Library Preparation (Illumina protocol, Part # 15044223 Rev. B) with the following modifications: (1) ITS2 primers (ITSD-forward: 5’-TCG TCG GCA GCG TCA GAT GTG TAT AAG AGA CAG GTG AAT TGC AGA ACT CCG TG-3’ 63 385 and ITS2-reverse: 5’-GTC TCG TGG GCT CGG AGA TGT GTA TAA GAG ACA GCC TCC GCT TAC TTA TAT GCT T-3’ 64) were used in place of the 16S primers. 2) PCR 1 annealing temperature was 52°C, PCR 1 was performed in triplicate, and PCR product was pooled prior to bead clean. 3) A 1:1.1 ratio of PCR product to SPRI beads was used for PCR 1 and PCR 2 clean up. Samples were sequenced on the Illumina MiSeq platform, which yielded 2×300 bp paired-end reads. Raw sequence data are available on the NCBI Sequence Read Archive under the BioProject accession PRJNA869694.

Extracts from n = 67 samples were prepared for 2b-RAD sequencing following the methods of Wang et al. (*75*). These samples were intended to cover a range of sites while focusing on the colonies for which survivorship was known. However, there was not enough tissue or DNA remaining to include all samples. Eight replicate samples were prepared to identify clones (none were found in this dataset). Samples were barcoded, multiplexed, and sequenced across two lanes of Illumina HiSeq 2500 at Tufts University Core Facility (TUCF). Raw reads were trimmed, deduplicated and quality filtered with FASTX TOOLKIT (http://hannonlab.cshl.edu/fastx_toolkit) and only reads with Phred scores >20 were maintained (-q 20 -p 100).

Quality-filtered reads were first mapped to a concatenated genome of four Symbiodiniaceae genera *Symbiodinium, Breviolum, Cladocopium*, and *Durusdinium* (*76–78*) via bowtie2 (*79*). Any reads that mapped successfully with a minimum end-to-end alignment score of − 22.2 were removed so that those left behind could be assumed to belong to the host. Remaining reads were then mapped to the *Porites lutea* genome (*80*). Genotyping and identification of single nucleotide polymorphisms (SNPs) was performed using ANGSD v0.921 (*81*). Standard filtering that was used across all analyses included loci present in at least 80% of individuals, minimum mapping quality score of 20, minimum quality score of 25 (unless no minimum allele frequency (MAF) filter was used in which case quality scores of 25 and 30 were used), strand bias p-value > 0.05, heterozygosity bias >0.05, removing all triallelic sites, removing reads having multiple best hits and lumped paralogs filter (see Supp. File 2 for proportion of missing data for each analysis).

### Lineage assignment

To detect population structure among corals from all sites, the program ADMIXTURE v. 1.3.0 (*82*) was used to find the optimal number of clusters (K) with the least cross validation error. SNPs were hard called using genotype likelihoods estimated by SAMtools with a SNP p-value < 0.05 (12,755 loci). Principal Coordinate Analyses (PCoAs; using 1-Pearson correlation) were performed using a covariance matrix based on single-read resampling calculated in ANGSD and admixture results were visualized using the K with the least cross validation error reported from ADMIXTURE and the most likely K based on the PCA. Samples were assigned to lineages based on the >0.85 assignment to a single lineage in ADMIXTURE and segregation along PC1 and PC2 axes in PCoA space.

Following lineage assignment using the 2b-RAD data, we used host contamination in the ITS2 metabarcoding data to further assign additional colonies to each lineage using a DNA barcoding approach. Sequence files generated with the intent of characterizing algal symbiont communities were run through the dada2 pipeline (*83*) in R using a reference database that included both Symbiodiniaceae and *Porites* ITS2 sequences (taken from Genbank; see Fig S4). Amplicon sequence variants (ASVs) matching *Porites* were isolated and the dominant ASV (for homozygous; at least 97% relative read abundance) or top two ASVs (for heterozygous; at least 40% relative abundance for second most abundant sequence) found in each coral were treated as DNA barcodes and used to assign lineages for samples not sequenced using 2b-RAD. For colonies that had ambiguous sequences or sequences that did not match any references (i.e., samples used for 2b-RAD), lineage assignment was not possible.

### Analyses of genetic divergence and demographics between lineages

BayeScan v. 2.1 (*84*) was used to identify a set of putatively neutral loci. The FST outlier method implemented in BayeScan identified outliers using 5000 iterations, 20 pilot runs with length 5000, and burn-in length of 50,000. We employed the default prior odds of neutrality (10) and a q-value cut-off of 0.50 after FDR correction for removing all putatively non-neutral loci. To determine genetic differentiation between lineages, ANGSD was used to calculate the site allele frequency (SAF) for each lineage using no MAF filter (363,736 loci) and then realSFS calculated the site frequency spectrum (SFS) for all possible pairwise comparisons. These SFSs were used as priors with the SAF to calculate global FST. Here, only weighted global FST values between lineages are reported. ANGSD was used to obtain 100 series of 5 block-bootstrapped SFS replicates, which were averaged to create 100 bootstrapped SFS for each lineage. SFS was polarized using the *P. lutea* genome as an ancestral reference. Multimodel inference in moments was used to fit two-population models (https://github.com/z0on/AFS-analysis-with-moments) and all unfolded models were run on 10 bootstrapped SFS and replicated six times. The best fit model was then selected based on lowest AIC value. Parameters (i.e., migration, epoch times, and effective population sizes (Ne)) for the best fit model were obtained by running the best fit model on 100 bootstrapped SFS and replicated six times. Additionally, we ran the unsupervised analysis StairwayPlot v2 (*85*) to one dimensional SFS as a second effort to reconstruct effective population sizes. For all demographic analyses we used a mutation rate 1.38e-9 (from the 0.138% per Ma substitution rate in Prada et al (*86*) calculated for the *Porites* genus) per base per year and generation time of 6 years. The generation time was calculated from the average reproductive age of *P. lutea* (8cm diameter; (*87*)) and average growth rate of 1.3 + 0.3 cm/year for *P. lobata* (*88*).

### Identifying genes under selection across lineages

Additional filtering of loci was conducted prior to outlier analyses, which included SNP p-value e− 5 (SNPs were hard called for this analysis) and MAF < 0.05 (4,956 loci). Our data were subset to include only two pairs of lineages for each comparison. The aim of this approach was to isolate outlier loci between the PKir-3 and versus both PKir-1 and PKir-2 to look for candidate genes that might explain the differential mortality outcomes. First, PCAdapt v. 4.3.3 (*89*) was used to determine the optimal K for all pairwise comparisons using a score plot displaying population structure. A K of 2 was selected for all pairwise comparisons between all lineage pairs and p-values were extracted from PC1, which separated each lineage pair. We performed an FDR correction on these p-values to create converted q-values, which were transformed using a BH correction to account for the multiple comparisons between lineages. A q-value of 0.05 was used as a cutoff for determining outlier loci and annotated genes (using the annotation file from (*90*)) 1 kb upstream or downstream of this outlier locus were reported.

### Analysis of algal symbiont communities

Symbiodiniaceae communities were inferred via ITS2 sequence data using SymPortal, implemented through the online portal (*50*). Analyses were conducted (and visualizations were produced) using both the ITS2 profile matrix and DIV matrix output directly from SymPortal (see below). In order to produce evolutionary trees for unifrac-based ordinations of the DIV matrix, sequences were aligned in Geneious and a NJ tree was produced. We used unifrac dissimilarity matrices (taken directly from SymPortal) to produce a NJ tree of *Cladocopium* ITS2 profiles.

### Statistical analyses

To test whether lineages were non-randomly distributed across the island and across the human disturbance gradient before the heatwave, we conducted Bayes Factor contingency tests. For geographic effects, we divided the island into four regions (North Lagoon Face n = 3 sites, Vaskess Bay/South Lagoon Face; n = 5 sites, Bay of Wrecks; n = 2 sites and North Shore; n = 3 sites), while we treated human disturbance as a continuous metric. We also tested whether algal symbiont communities differed across lineages before the heatwave, using a PERMANOVA on the DIV matrix output from SymPortal and we visualized ordinations (Fig 3a,b) using only sequence reads from the *Cladocopium* C15 clade. We tested for effects of coral lineage and human disturbance on coral survival, by conducting a binomial logistic regression with these two variables and an interaction term between them. We tested for differences in categorical bleaching status across lineages both early and late in the heatwave using ordinal logistic regressions. These analyses were run using the following packages in R: bayesfactor, dada2, phyloseq, tidyverse, vegan, and vgam.

## Acknowledgments

We are grateful to the Government of Kiribati, and the people of Kiritimati for their support of our research over several years. We acknowledge with respect that the University of Victoria stands on the traditional territory of the Lekwangen speaking peoples, including the Songhees, Esquimalt and □ SÁNEĆ nations whose relationships with the land continue to this day. We also acknowledge that research at the Boston University was performed on the ancestral land of the Pawtucket, Massachusett, and Naumkeag tribes. Thanks to J. Davidson for logistical and lab support, A. Eggersfor for molecular sequencing support, B. Koop for providing laboratory space and equipment. We thank also B. Hume for support analyzing metabarcode data in SymPortal. Analysis of genomic data was made possible through BU’s Shared Computing Cluster.

## Funding

NSERC Postdoctoral Fellowship

NSERC Discovery Grant

NOAA Climate and Global Change Postdoctoral Fellowship Program #NA18NWS4620043B

NSF RAPID (OCE-1446402)

David and Lucile Packard Foundation

Rufford Maurice Laing Foundation

Pew Fellowship in Marine Conservation

NSF OCE-1358699

NSF OCE-1851392

Boston University (start-up funding)

## Author contributions

Conceptualization: SS, JF, DCC, SWD, RC, ACB, JKB

Methodology: SS, JF, DCC, SWD, RC, ACB, JKB

Visualization: SS, JF Supervision: SWD, JKB

Writing—original draft: SS, JF

Writing—review & editing: SS, JF, DCC, SWD, RC, ACB, JKB

### Data and materials availability

All data and code will be made available on Zenodo upon acceptance.

## References

1. A. Rammig, M. D. Mahecha, Ecology: Ecosystem responses to climate extremes. Nature. 527, 315–316 (2015).

2. M. D. Smith, The ecological role of climate extremes: current understanding and future prospects. Journal of Ecology. 99, 651–655 (2011).

3. S. Legg, Climate Change 2021-the Physical Science basis. International Panel on Climate Change. 49, 44–45 (2021).

4. T. Wernberg, S. Bennett, R. C. Babcock, T. de Bettignies, K. Cure, M. Depczynski, F. Dufois, J. Fromont, C. J. Fulton, R. K. Hovey, E. S. Harvey, T. H. Holmes, G. A. Kendrick, B. Radford, J. Santana-Garcon, B. J. Saunders, D. A. Smale, M. S. Thomsen, C. A. Tuckett, F. Tuya, M. A. Vanderklift, S. Wilson, Climate-driven regime shift of a temperate marine ecosystem. Science. 353, 169–172 (2016).

5. D. A. Smale, T. Wernberg, E. C. J. Oliver, M. Thomsen, B. P. Harvey, S. C. Straub, M. T. Burrows, L. V. Alexander, J. A. Benthuysen, M. G. Donat, M. Feng, A. J. Hobday, N. J. Holbrook, S. E. Perkins-Kirkpatrick, H. A. Scannell, A. S. Gupta, B. L. Payne, P. J. Moore, Marine heatwaves threaten global biodiversity and the provision of ecosystem services. Nature Climate Change, 1 (2019).

6. E. I. A. y Juárez, E. A. Ellis, E. Rodríguez-Luna, Quantifying the severity of hurricanes on extinction probabilities of a primate population: Insights into “Island” extirpations. American Journal of Primatology. 77, 786–800 (2015).

7. M. Romero-Torres, A. Acosta, A. M. Palacio-Castro, E. A. Treml, F. A. Zapata, D. A. Paz-García, J. W. Porter, Coral reef resilience to thermal stress in the Eastern Tropical Pacific. Global Change Biology. 26, 3880–3890 (2020).

8. H. Tanaka, M. Yasuhara, J. T. Carlton, Transoceanic transport of living marine Ostracoda (Crustacea) on tsunami debris from the 2011 Great East Japan Earthquake. Aquatic Invasions. 13 (2018).

9. P. R. Grant, B. R. Grant, R. B. Huey, M. T. J. Johnson, A. H. Knoll, J. Schmitt, Evolution caused by extreme events. Philosophical Transactions of the Royal Society B: Biological Sciences. 372, 20160146 (2017).

10. M. A. Coleman, T. Wernberg, The Silver Lining of Extreme Events. Trends in Ecology & Evolution (2020), doi:10.1016/j.tree.2020.08.013.

11. T. C. LaJeunesse, R. Smith, M. Walther, J. Pinzón, D. T. Pettay, M. McGinley, M. Aschaffenburg, P. Medina-Rosas, A. L. Cupul-Magaña, A. L. Pérez, H. Reyes-Bonilla, M. E. Warner, Host–symbiont recombination versus natural selection in the response of coral–dinoflagellate symbioses to environmental disturbance. Proceedings of the Royal Society B: Biological Sciences. 277, 2925–2934 (2010).

12. C. Gurgel, O. Camacho, A. Minne, T. Wernberg, M. Coleman, Marine heatwave drives cryptic loss of genetic diversity in underwater forests. Current Biology. 30 (2020).

13. A. G. Little, D. N. Fisher, T. W. Schoener, J. N. Pruitt, Population differences in aggression are shaped by tropical cyclone-induced selection. Nat Ecol Evol. 3, 1294–1297 (2019).

14. S. C. Campbell-Staton, Z. A. Cheviron, N. Rochette, J. Catchen, J. B. Losos, S. V. Edwards, Winter storms drive rapid phenotypic, regulatory, and genomic shifts in the green anole lizard. Science. 357, 495–498 (2017).

15. D. Lirman, S. Schopmeyer, D. Manzello, L. J. Gramer, W. F. Precht, F. Muller-Karger, K. Banks, B. Barnes, E. Bartels, A. Bourque, J. Byrne, S. Donahue, J. Duquesnel, L. Fisher, D. Gilliam, J. Hendee, M. Johnson, K. Maxwell, E. McDevitt, J. Monty, D. Rueda, R. Ruzicka, S. Thanner, Severe 2010 Cold-Water Event Caused Unprecedented Mortality to Corals of the Florida Reef Tract and Reversed Previous Survivorship Patterns. PLOS ONE. 6, e23047 (2011).

16. S. U. Pauls, C. Nowak, M. Bálint, M. Pfenninger, The impact of global climate change on genetic diversity within populations and species. Molecular Ecology. 22, 925–946 (2013).

17. T. P. Hughes, K. D. Anderson, S. R. Connolly, S. F. Heron, J. T. Kerry, J. M. Lough, A. H. Baird, J. K. Baum, M. L. Berumen, T. C. Bridge, D. C. Claar, C. M. Eakin, J. P. Gilmour, N. A. J. Graham, H. Harrison, J.-P. A. Hobbs, A. S. Hoey, M. Hoogenboom, R. J. Lowe, M. T. McCulloch, J. M. Pandolfi, M. Pratchett, V. Schoepf, G. Torda, S. K. Wilson, Spatial and temporal patterns of mass bleaching of corals in the Anthropocene. Science. 359, 80–83 (2018).

18. A. J. Hobday, L. V. Alexander, S. E. Perkins, D. A. Smale, S. C. Straub, E. C. Oliver, J. A. Benthuysen, M. T. Burrows, M. G. Donat, M. Feng, A hierarchical approach to defining marine heatwaves. Progress in Oceanography. 141, 227–238 (2016).

19. T. L. Frölicher, E. M. Fischer, N. Gruber, Marine heatwaves under global warming. Nature. 560, 360–364 (2018).

20. J. M. T. Magel, S. A. Dimoff, J. K. Baum, Direct and indirect effects of climate change-amplified pulse heat stress events on coral reef fish communities. Ecological Applications. 30, e02124 (2020).

21. S. P. Brady, D. I. Bolnick, A. L. Angert, A. Gonzalez, R. D. H. Barrett, E. Crispo, A. M. Derry, C. G. Eckert, D. J. Fraser, G. F. Fussmann, F. Guichard, T. Lamy, A. G. McAdam, A. E. M. Newman, A. Paccard, G. Rolshausen, A. M. Simons, A. P. Hendry, Causes of maladaptation. Evolutionary Applications. 12, 1229– 1242 (2019).

22. S. C. Burgess, E. C. Johnston, A. S. Wyatt, J. J. Leichter, P. J. Edmunds, Response diversity in corals: hidden differences in bleaching mortality among cryptic Pocillopora species. Ecology. 102, e03324 (2021).

23. H.-O. Pörtner, D. C. Roberts, V. Masson-Delmotte, P. Zhai, M. Tignor, E. Poloczanska, K. Mintenbeck, M. Nicolai, A. Okem, J. Petzold, IPCC special report on the ocean and cryosphere in a changing climate. IPCC Intergovernmental Panel on Climate Change: Geneva, Switzerland. (2019).

24. R. W. Buddemeier, D. G. Fautin, Coral bleaching as an adaptive mechanism. Bioscience. 43, 320–326 (1993).

25. T. C. LaJeunesse, J. E. Parkinson, P. W. Gabrielson, H. J. Jeong, J. D. Reimer, C. R. Voolstra, S. R. Santos, Systematic revision of Symbiodiniaceae highlights the antiquity and diversity of coral endosymbionts. Current Biology. 28, 2570-2580.e6 (2018).

26. P. W. Glynn, Coral reef bleaching: facts, hypotheses and implications. Global Change Biology. 2, 495–509 (1996).

27. J. P. G. Spurgeon, The economic valuation of coral reefs. Marine Pollution Bulletin. 24, 529–536 (1992).

28. M. J. H. van Oppen, J. K. Oliver, H. M. Putnam, R. D. Gates, Building coral reef resilience through assisted evolution. Proc. Natl. Acad. Sci. U.S.A. 112, 2307–2313 (2015).

29. A. C. Baker, C. J. Starger, T. R. McClanahan, P. W. Glynn, Corals’ adaptive response to climate change. Nature. 430, 741–741 (2004).

30. R. Cunning, R. N. Silverstein, A. C. Baker, Investigating the causes and consequences of symbiont shuffling in a multi-partner reef coral symbiosis under environmental change. Proc. R. Soc. B. 282, 20141725 (2015).

31. M. Stat, R. D. Gates, Clade D Symbiodinium in scleractinian corals: A “nugget” of hope, a selfish opportunist, an ominous Sign, or all of the above? Journal of Marine Biology. 2011, e730715 (2011).

32. K. M. Quigley, B. L. Willis, L. K. Bay, Heritability of the Symbiodinium community in vertically-and horizontally-transmitting broadcast spawning corals. Sci Rep. 7, 8219 (2017).

33. N. H. Rose, R. A. Bay, M. K. Morikawa, L. Thomas, E. A. Sheets, S. R. Palumbi, Genomic analysis of distinct bleaching tolerances among cryptic coral species. Proc. R. Soc. B-Biol. Sci. 288, 20210678 (2021).

34. D. C. Claar, S. Starko, K. L. Tietjen, H. E. Epstein, R. Cunning, K. M. Cobb, A. C. Baker, R. D. Gates, J. K. Baum, Dynamic symbioses reveal pathways to coral survival through prolonged heatwaves. Nature Communications. 11, 6097 (2020).

35. E. C. Johnston, R. Cunning, S. C. Burgess, Cophylogeny and specificity between cryptic coral species (Pocillopora spp.) at Mo’orea and their symbionts (Symbiodiniaceae) (2022), p. 2022.03.02.482706,, doi:10.1101/2022.03.02.482706.

36. M. J. van Oppen, P. Bongaerts, P. Frade, L. M. Peplow, S. E. Boyd, H. T. Nim, L. K. Bay, Adaptation to reef habitats through selection on the coral animal and its associated microbiome. Molecular ecology. 27, 2956– 2971 (2018).

37. S. V. Vollmer, S. R. Palumbi, Hybridization and the Evolution of Reef Coral Diversity. Science. 296, 2023– 2025 (2002).

38. J. T. Ladner, S. R. Palumbi, Extensive sympatry, cryptic diversity and introgression throughout the geographic distribution of two coral species complexes. Molecular Ecology. 21, 2224–2238 (2012).

39. K. R. Hind, S. Starko, J. M. Burt, M. A. Lemay, A. K. Salomon, P. T. Martone, Trophic control of cryptic coralline algal diversity. PNAS. 116, 15080–15085 (2019).

40. M. J. Brasier, H. Wiklund, L. Neal, R. Jeffreys, K. Linse, H. Ruhl, A. G. Glover, DNA barcoding uncovers cryptic diversity in 50% of deep-sea Antarctic polychaetes. Royal Society Open Science. 3, 160432 (2016).

41. Z. H. Forsman, D. J. Barshis, C. L. Hunter, R. J. Toonen, Shape-shifting corals: Molecular markers show morphology is evolutionarily plastic in Porites. BMC Evolutionary Biology. 9, 45 (2009).

42. Z. H. Forsman, R. Ritson-Williams, K. H. Tisthammer, I. S. S. Knapp, R. J. Toonen, Host-symbiont coevolution, cryptic structure, and bleaching susceptibility, in a coral species complex (Scleractinia; Poritidae). Scientific reports. 10, 1–12 (2020).

43. J. E. Fifer, N. Yasuda, T. Yamakita, C. B. Bove, S. W. Davies, Genetic divergence and range expansion in a western North Pacific coral. Science of The Total Environment. 813, 152423 (2022).

44. M. GómezΔCorrales, C. Prada, Cryptic lineages respond differently to coral bleaching. Molecular Ecology. 29, 4265–4273 (2020).

45. C. M. Eakin, H. P. Sweatman, R. E. Brainard, The 2014–2017 global-scale coral bleaching event: insights and impacts. Coral Reefs. 38, 539–545 (2019).

46. T. P. Hughes, J. T. Kerry, M. Álvarez-Noriega, J. G. Álvarez-Romero, K. D. Anderson, A. H. Baird, R. C. Babcock, M. Beger, D. R. Bellwood, R. Berkelmans, Global warming and recurrent mass bleaching of corals. Nature. 543, 373–377 (2017).

47. T. D. Ainsworth, S. F. Heron, J. C. Ortiz, P. J. Mumby, A. Grech, D. Ogawa, C. M. Eakin, W. Leggat, Climate change disables coral bleaching protection on the Great Barrier Reef. Science. 352, 338–342 (2016).

48. J. K. Baum, D. C. Claar, K. L. Tietjen, J. M. T. Magel, D. G. Maucieri, K. M. Cobb, J. M. McDevitt-Irwin, Transformation of coral communities subjected to an unprecedented heatwave is modulated by local disturbance (2022), p. 2022.05.10.491220,, doi:10.1101/2022.05.10.491220.

49. T. I. Terraneo, F. Benzoni, R. Arrigoni, A. H. Baird, K. G. Mariappan, Z. H. Forsman, M. K. Wooster, J. Bouwmeester, A. Marshell, M. L. Berumen, Phylogenomics of Porites from the Arabian Peninsula. Molecular Phylogenetics and Evolution. 161, 107173 (2021).

50. B. C. Hume, E. G. Smith, M. Ziegler, H. J. Warrington, J. A. Burt, T. C. LaJeunesse, J. Wiedenmann, C. R. Voolstra, SymPortal: A novel analytical framework and platform for coral algal symbiont nextΔgeneration sequencing ITS2 profiling. Molecular Ecology Resources. 19, 1063–1080 (2019).

51. C. D. Kenkel, L. K. Bay, Exploring mechanisms that affect coral cooperation: symbiont transmission mode, cell density and community composition. PeerJ. 6, e6047 (2018).

52. A. H. Moeller, A. Caro-Quintero, D. Mjungu, A. V. Georgiev, E. V. Lonsdorf, M. N. Muller, A. E. Pusey, M. Peeters, B. H. Hahn, H. Ochman, Cospeciation of gut microbiota with hominids. Science. 353, 380–382 (2016).

53. A. H. Moeller, T. A. Suzuki, M. Phifer-Rixey, M. W. Nachman, Transmission modes of the mammalian gut microbiota. Science. 362, 453–457 (2018).

54. A. Hayward, R. Poulin, S. Nakagawa, A broadscale analysis of hostΔsymbiont cophylogeny reveals the drivers of phylogenetic congruence. Ecology Letters. 24, 1681–1696 (2021).

55. A. C. Baker, Flexibility and specificity in coral-algal symbiosis: diversity, ecology, and biogeography of Symbiodinium. Annual Review of Ecology, Evolution, and Systematics. 34, 661–689 (2003).

56. K. D. Hoadley, D. T. Pettay, A. Lewis, D. Wham, C. Grasso, R. Smith, D. W. Kemp, T. LaJeunesse, M. E. Warner, Different functional traits among closely related algal symbionts dictate stress endurance for vital Indo-Pacific reef-building corals. Glob Chang Biol. 27, 5295–5309 (2021).

57. E. J. Howells, V. H. Beltran, N. W. Larsen, L. K. Bay, B. L. Willis, M. J. H. van Oppen, Coral thermal tolerance shaped by local adaptation of photosymbionts. Nature Climate Change. 2, 116–120 (2012).

58. S. A. Fay, M. X. Weber, The Occurrence of Mixed Infections of Symbiodinium (Dinoflagellata) within Individual Hosts. Journal of Phycology. 48, 1306–1316 (2012).

59. Y. T. R. Tan, B. J. Wainwright, L. Afiq-Rosli, Y. C. A. Ip, J. N. Lee, N. T. H. Nguyen, S. B. Pointing, D. Huang, Endosymbiont diversity and community structure in Porites lutea from Southeast Asia are driven by a suite of environmental variables. Symbiosis. 80, 269–277 (2020).

60. A. C. Baker, T. R. McClanahan, C. J. Starger, R. K. Boonstra, Long-term monitoring of algal symbiont communities in corals reveals stability is taxon dependent and driven by site-specific thermal regime. Marine Ecology Progress Series. 479, 85–97 (2013).

61. K. M. Quigley, P. A. Warner, L. K. Bay, B. L. Willis, Unexpected mixed-mode transmission and moderate genetic regulation of Symbiodinium communities in a brooding coral. Heredity (Edinb). 121, 524–536 (2018).

62. T. I. Terraneo, M. Fusi, B. C. C. Hume, R. Arrigoni, C. R. Voolstra, F. Benzoni, Z. H. Forsman, M. L. Berumen, Environmental latitudinal gradients and host-specificity shape Symbiodiniaceae distribution in Red Sea Porites corals. Journal of Biogeography. 46, 2323–2335 (2019).

63. M. Ziegler, C. M. Roder, C. Büchel, C. R. Voolstra, Mesophotic coral depth acclimatization is a function of host-specific symbiont physiology. Frontiers in Marine Science. (2015) (available at https://www.frontiersin.org/article/10.3389/fmars.2015.00004).

64. M. T. Connelly, C. J. McRae, P.-J. Liu, N. Traylor-Knowles, Lipopolysaccharide treatment stimulates Pocillopora coral genotype-specific immune responses but does not alter coral-associated bacteria communities. Developmental & Comparative Immunology. 109, 103717 (2020).

65. D. Bickford, D. J. Lohman, N. S. Sodhi, P. K. L. Ng, R. Meier, K. Winker, K. K. Ingram, I. Das, Cryptic species as a window on diversity and conservation. Trends in Ecology & Evolution. 22, 148–155 (2007).

66. L. Rüber, J. L. Van Tassell, R. Zardoya, S. Karl, Rapid speciation and ecological divergence in the american seven-spined gobies (gobiidae, Gobiosomatini) inferred from a molecular phylogeny. Evolution. 57, 1584– 1598 (2003).

67. D. Schluter, Ecological causes of adaptive radiation. The American Naturalist. 148, S40–S64 (1996).

68. S. Starko, K. Demes, C. Neufeld, P. Martone, Convergent evoliution of niche structure in Northeast Pacific kelp forests. Functional Ecology (2020), doi:https://doi.org/10.1111/1365-2435.13621.

69. D. Ackerly, Conservatism and diversification of plant functional traits: Evolutionary rates versus phylogenetic signal. PNAS. 106, 19699–19706 (2009).

70. N. N. Winchester, R. A. Ring, CENTINELAN EXTINCTIONS: EXTIRPATION OF NORTHERN TEMPERATE OLD-GROWTH RAINFOREST ARTHROPOD COMMUNITIES. Selbyana. 17, 50–57 (1996).

71. O. Morate, Population and Housing Census Volume 1: Management Report and Basic Tables, (2015).

72. M. S. Watson, D. C. Claar, J. K. Baum, Subsistence in isolation: Fishing dependence and perceptions of change on Kiritimati, the world’s largest atoll. Ocean & coastal management. 123, 1–8 (2016).

73. M. Stat, W. K. W. Loh, T. C. LaJeunesse, O. Hoegh-Guldberg, D. A. Carter, Stability of coral–endosymbiont associations during and after a thermal stress event in the southern Great Barrier Reef. Coral Reefs. 28, 709– 713 (2009).

74. I. B. Baums, C. R. Hughes, M. E. Hellberg, Mendelian microsatellite loci for the Caribbean coral Acropora palmata. Marine Ecology Progress Series. 288, 115–127 (2005).

75. S. Wang, E. Meyer, J. K. McKay, M. V. Matz, 2b-RAD: a simple and flexible method for genome-wide genotyping. Nat Methods. 9, 808–810 (2012).

76. M. Aranda, Y. Li, Y. J. Liew, S. Baumgarten, O. Simakov, M. C. Wilson, J. Piel, H. Ashoor, S. Bougouffa, V. B. Bajic, T. Ryu, T. Ravasi, T. Bayer, G. Micklem, H. Kim, J. Bhak, T. C. LaJeunesse, C. R. Voolstra, Genomes of coral dinoflagellate symbionts highlight evolutionary adaptations conducive to a symbiotic lifestyle. Sci Rep. 6, 39734 (2016).

77. H. Liu, T. G. Stephens, R. A. González-Pech, V. H. Beltran, B. Lapeyre, P. Bongaerts, I. Cooke, M. Aranda, D. G. Bourne, S. Forêt, D. J. Miller, M. J. H. van Oppen, C. R. Voolstra, M. A. Ragan, C. X. Chan, Symbiodinium genomes reveal adaptive evolution of functions related to coral-dinoflagellate symbiosis. Commun Biol. 1, 1–11 (2018).

78. E. Shoguchi, C. Shinzato, T. Kawashima, F. Gyoja, S. Mungpakdee, R. Koyanagi, T. Takeuchi, K. Hisata, M. Tanaka, M. Fujiwara, M. Hamada, A. Seidi, M. Fujie, T. Usami, H. Goto, S. Yamasaki, N. Arakaki, Y. Suzuki, S. Sugano, A. Toyoda, Y. Kuroki, A. Fujiyama, M. Medina, M. A. Coffroth, D. Bhattacharya, N. Satoh, Draft Assembly of the Symbiodinium minutum Nuclear Genome Reveals Dinoflagellate Gene Structure. Current Biology. 23, 1399–1408 (2013).

79. B. Langmead, S. L. Salzberg, Fast gapped-read alignment with Bowtie 2. Nat Methods. 9, 357–359 (2012).

80. Y. J. Liew, M. Aranda, C. R. Voolstra, Reefgenomics.Org - a repository for marine genomics data. Database. 2016, baw152 (2016).

81. T. S. Korneliussen, A. Albrechtsen, R. Nielsen, ANGSD: analysis of next generation sequencing data. BMC bioinformatics. 15, 1–13 (2014).

82. D. H. Alexander, S. S. Shringarpure, J. Novembre, K. Lange, Admixture 1.3 software manual. Los Angeles: UCLA Human Genetics Software Distribution (2015).

83. B. J. Callahan, P. J. McMurdie, M. J. Rosen, A. W. Han, A. J. A. Johnson, S. P. Holmes, DADA2: High-resolution sample inference from Illumina amplicon data. Nature methods. 13, 581–583 (2016).

84. M. Foll, BayeScan v2. 1 user manual. Ecology. (2012).

85. X. Liu, Y.-X. Fu, Stairway Plot 2: demographic history inference with folded SNP frequency spectra.Genome Biology. 21, 280 (2020).

86. C. Prada, M. B. DeBiasse, J. E. Neigel, B. Yednock, J. L. Stake, Z. H. Forsman, I. B. Baums, M. E. Hellberg, Genetic species delineation among branching Caribbean Porites corals. Coral Reefs. 33, 1019–1030 (2014).

87. V. J. Harriott, Reproductive ecology of four scleratinian species at Lizard Island, Great Barrier Reef. Coral Reefs. 2, 9–18 (1983).

88. J. Pätzold, Growth rhythms recorded in stable isotopes and density bands in the reef coral Porites lobata (Cebu, Philippines). Coral Reefs. 3, 87–90 (1984).

89. K. Luu, E. Bazin, M. G. B. Blum, pcadapt: an R package to perform genome scans for selection based on principal component analysis. Molecular Ecology Resources. 17, 67–77 (2017).

90. H. Rivera, A. Cohen, J. Thompson, I. Baums, M. Fox, K. Meyer, Palau’s warmest reefs harbor a thermally tolerant coral lineage that thrives across different habitats (2022).

