## Supplementary Material for "Marine heatwaves threaten cryptic coral diversity and erode associations amongst coevolving partners"

**This PDF file includes:**

Figs. S1 to S8
Tables S1 to S5

**Other Supplementary Materials for this manuscript include the following:**

Supp File 1 to 2

Table S1. Global $F_{st}$ between lineage pairs.

| | *PKir* - 1 | *PKir* - 2 | *PKir* - 3 |
|---|---|---|---|
| *PKir* - 1 | - | 0.263 | 0.361 |
| *PKir* - 2 | 0.263 | - | 0.236 |
| *PKir* - 3 | 0.361 | 0.326 | - |

Table S2. NCBI accession numbers for *Porites* sequences used to assign lineage.

| ID IN THIS STUDY | SEQUENCE ID | GENBANK ACCESSION | LINEAGE |
|---|---|---|---|
| **ASV2** | PKir1-1 | *TBD* | PKir-1 |
| **ASV15** | PKir1-2 | *TBD* | PKir-1 |
| **ASV3** | PKir1-3 | *TBD* | PKir-1 |
| **ASV21** | PKir1-4 | *TBD* | PKir-1 |
| **ASV26** | PKir1-5 | *TBD* | PKir-1 |
| **ASV5** | PKir1-6 | *TBD* | PKir-1 |
| **ASV79** | PKir1-7 | *TBD* | PKir-1 |
| **ASV14** | PKir1-8 | *TBD* | PKir-1 |
| **ASV9** | PKir12-1 | *TBD* | PKir-1, PKir-2 |
| **ASV31** | PKir12-2 | *TBD* | PKir-1, PKir-2 |
| **ASV16** | PKir2-1 | *TBD* | PKir-2 |
| **ASV7** | PKir2-2 | *TBD* | PKir-2 |
| **ASV20** | PKir2-3 | *TBD* | PKir-2 |
| **ASV25** | PKir3-1 | *TBD* | PKir-3 |
| **ASV27** | PKir3-2 | *TBD* | PKir-3 |
| **ASV59** | PKir3-3 | *TBD* | PKir-3 |
| **ASV87** | PKir3-4 | *TBD* | PKir-3 |
| **ASV43** | PKir3-5 | *TBD* | PKir-3 |
| **ASV127** | PKir3-6 | *TBD* | PKir-3 |
| **ASV81** | PKir3-7 | *TBD* | PKir-3 |
| **ASV156** | PKir3-8 | *TBD* | PKir-3 |
| **ASV74** | PKir3-9 | *TBD* | PKir-3 |
| **ASV17** | PKir3-10 | *TBD* | PKir-3 |

Table S3. Summary of sample sizes by technique successfully used to characterize host or symbiont genotypes

| Technique | Sample size |
|---|---|
| **2bRAD + symbiont and host ITS2 sequences** | n = 64 |
| **2bRAD + symbiont ITS2 sequences only** | n = 3 |
| **Symbiont and host ITS2 sequences only** | n = 229 |
| **Host ITS2 sequences only** | n = 8 |
| **Symbiont ITS2 sequences only** | n = 1 |
| **Total number of colonies** | n = 305 |

Table S4. Summary of sample sizes for tracked colonies of known survival status (as of 2017).

| | PKir-1 | | PKir-2 | | PKir-3 | | Unassigned | | Total |
|---|---|---|---|---|---|---|---|---|---|
| Site | Survived | Died | Survived | Died | Survived | Died | Survived | Died | |
| VH1 | 0 | 1 | 0 | 1 | 0 | 3 | 0 | 0 | 5 |
| VH2 | 0 | 1 | 0 | 1 | 0 | 0 | 0 | 0 | 2 |
| VH3 | 0 | 5 | 0 | 5 | 1 | 1 | 0 | 1 | 13 |
| M1 | 2 | 1 | 1 | 0 | 0 | 3 | 0 | 0 | 7 |
| M2 | 2 | 4 | 1 | 0 | 0 | 2 | 0 | 0 | 9 |
| M3 | 3 | 1 | 0 | 1 | 1 | 2 | 0 | 0 | 8 |
| M5 | 2 | 1 | 2 | 0 | 0 | 0 | 0 | 0 | 5 |
| L1 | 1 | 0 | 0 | 0 | 1 | 1 | 0 | 0 | 3 |
| L2 | 0 | 0 | 2 | 0 | 0 | 0 | 0 | 0 | 2 |
| VL1 | 0 | 2 | 4 | 1 | 0 | 1 | 1 | 0 | 9 |
| VL2 | 0 | 1 | 4 | 0 | 0 | 2 | 0 | 0 | 7 |
| VL3 | 4 | 1 | 2 | 0 | 0 | 2 | 0 | 0 | 9 |
| Total | 14 | 18 | 16 | 9 | 3 | 17 | 1 | 1 | 79 |

Table S5. Summary of sample sizes by site for samples taken late during in the heatwave (2016 timepoint). Note that these include colonies for which either symbiont or host sequences were recovered (not necessarily both).

| Site | PKir-1 | PKir-2 | PKir-3 | Unassigned |
|---|---|---|---|---|
| VH1 | 1 | 2 | 1 | 0 |
| VH2 | 1 | 1 | 0 | 0 |
| VH3 | 1 | 1 | 0 | 0 |
| H1 | 7 | 2 | 0 | 3 |
| M1 | 4 | 2 | 1 | 0 |
| M2 | 7 | 1 | 0 | 0 |
| M3 | 4 | 2 | 1 | 0 |
| M12 | 4 | 2 | 1 | 0 |
| L2 | 1 | 3 | 0 | 0 |
| L4 | 3 | 1 | 0 | 0 |
| VL1 | 0 | 6 | 0 | 2 |
| VL3 | 10 | 4 | 3 | 1 |
| VL4 | 11 | 2 | 0 | 0 |

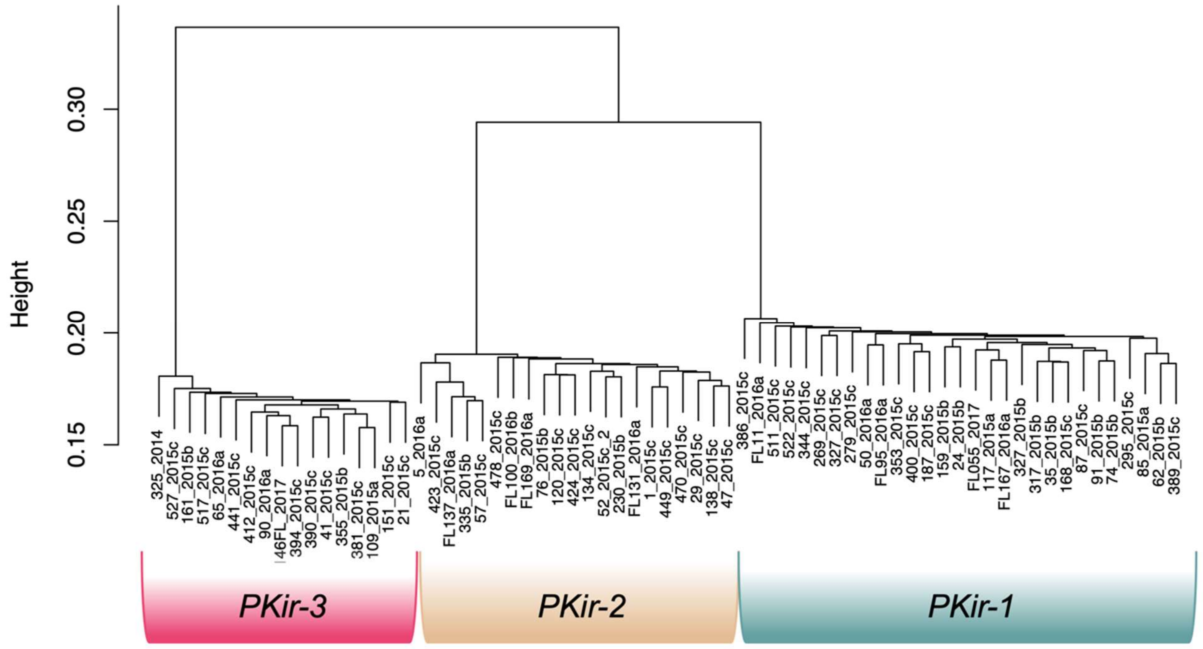


**Fig S1. Dendrogram showing relationships between colonies across and within lineages based on 2bRAD data.**

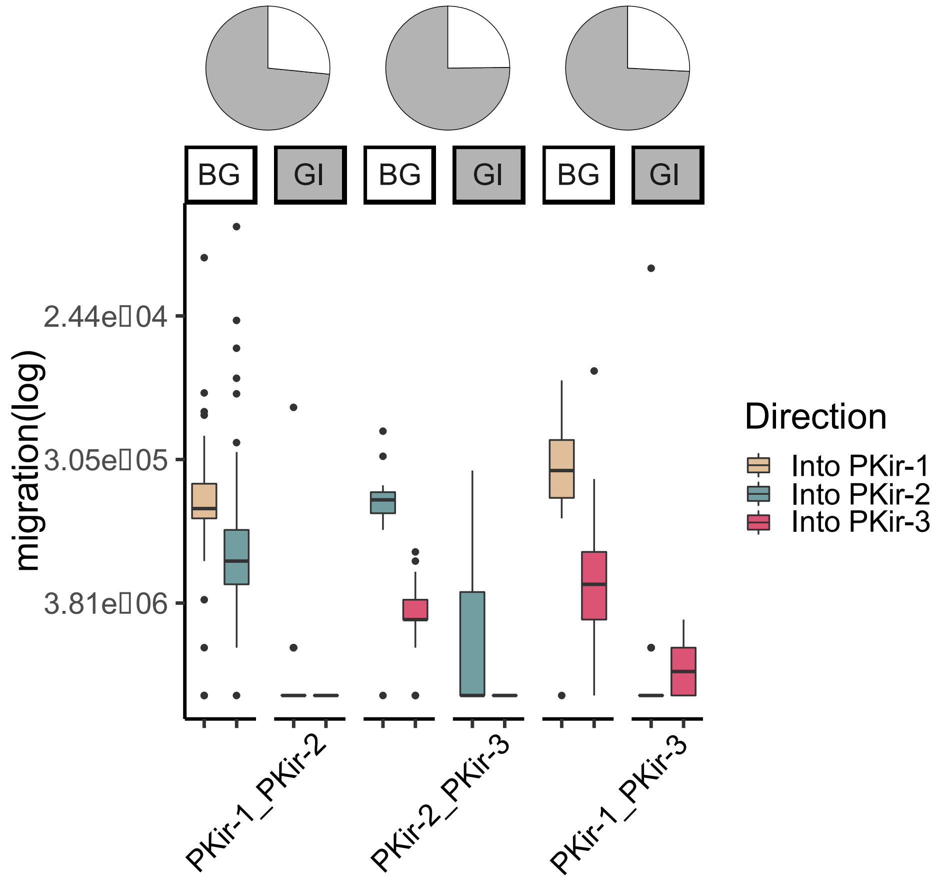


**Fig S2. Bootstrapped rates of gene flow between lineage pairs.** GI represents genomic islands of decreased gene flow relative to BG, the remainder of the genome. Note these GIs make up a majority of the genome (percentage shown in pie-chart).

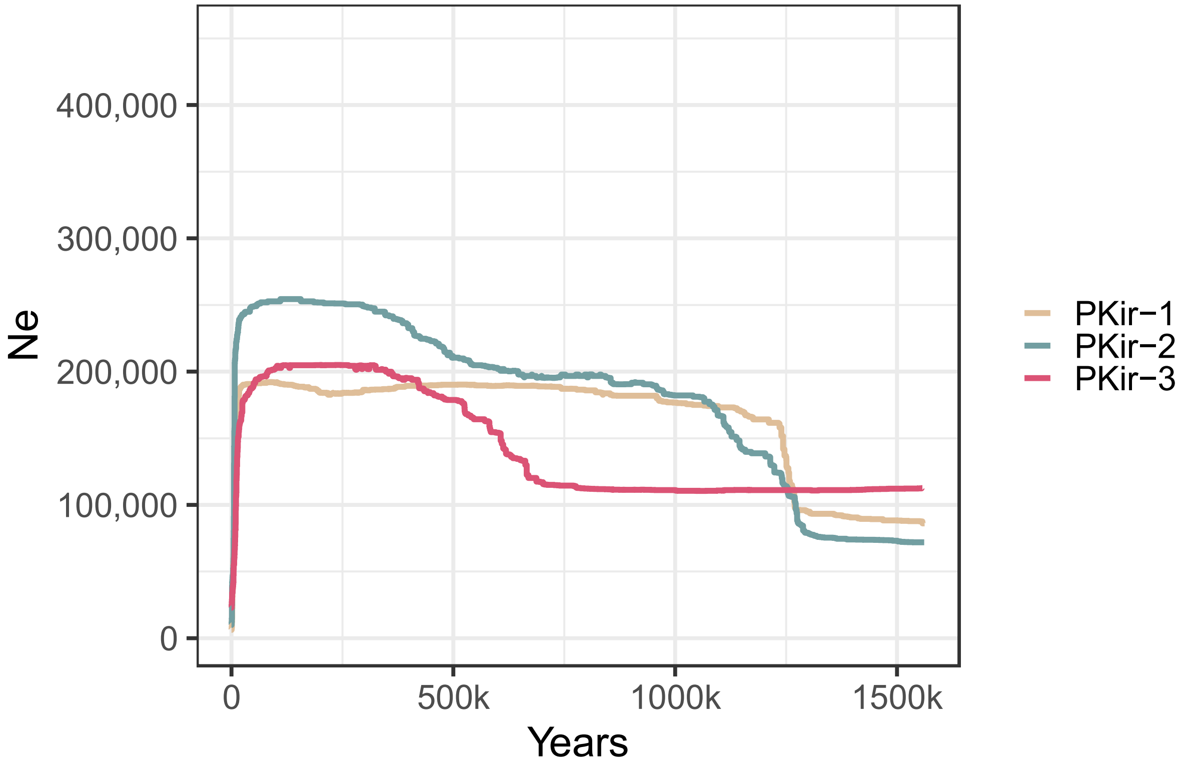


**Fig S3. Reconstruction of effective population size ($N_e$) over the past 1.5 million years in all three lineages of *Porites*.**

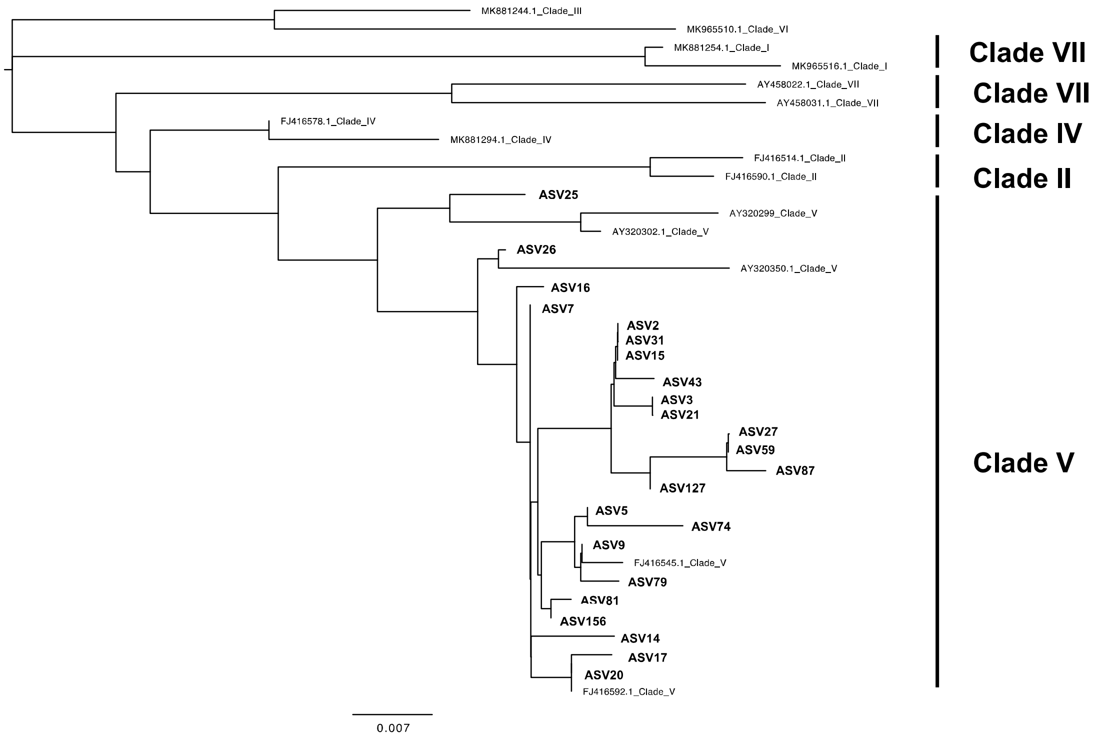


**Fig S4. Dominant ITS2 sequence variants from colonies in this study, compared to reference sequences from *Porites* clades I – VII.** Sequences from this study are shown in bold and labelled as ASVs.

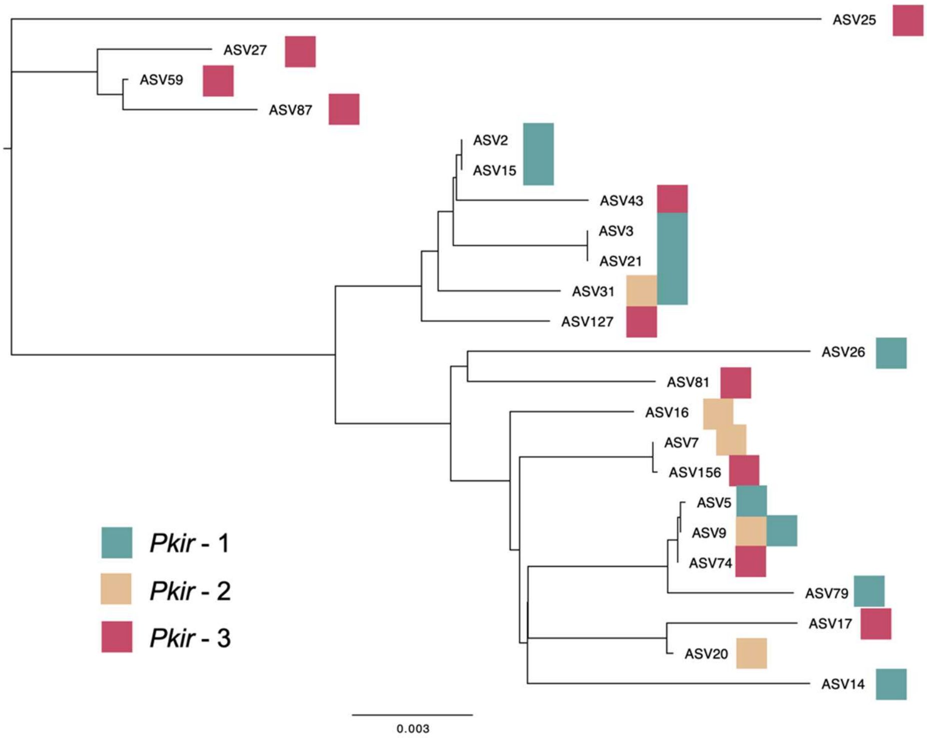


**Fig S5. Neighbour-joining tree of the ITS2 sequence variants used to assign colonies to lineage.** Each tip represents a sequence variant and the colour indicates the lineage(s) found to be associated with that particular variant. The tree was produced in Geneious Prime using a Tamura-Nei distance model. Sequence variants are from all colonies that were sequenced with both RADSeq and ITS2 barcoding (n = 65 colonies).

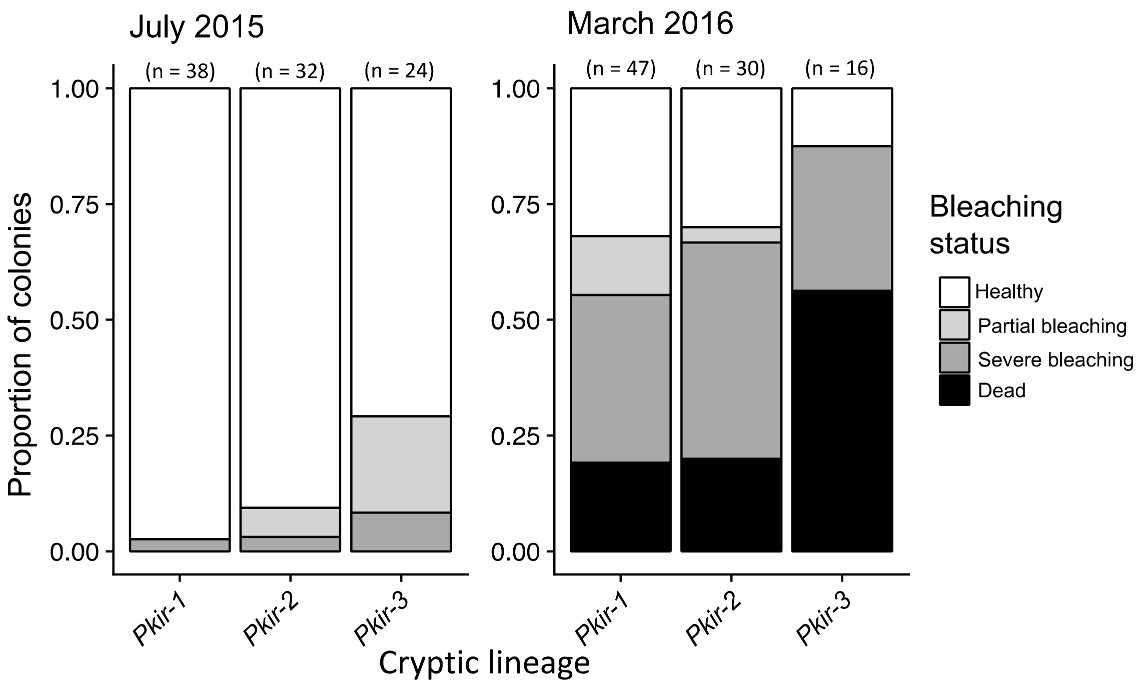


**Fig S6. Bleaching status of colonies early (July 2015) and late (March 2016) in the heatwave.** Shading indicates the degree of bleaching experienced by a particular colony at each timepoint.

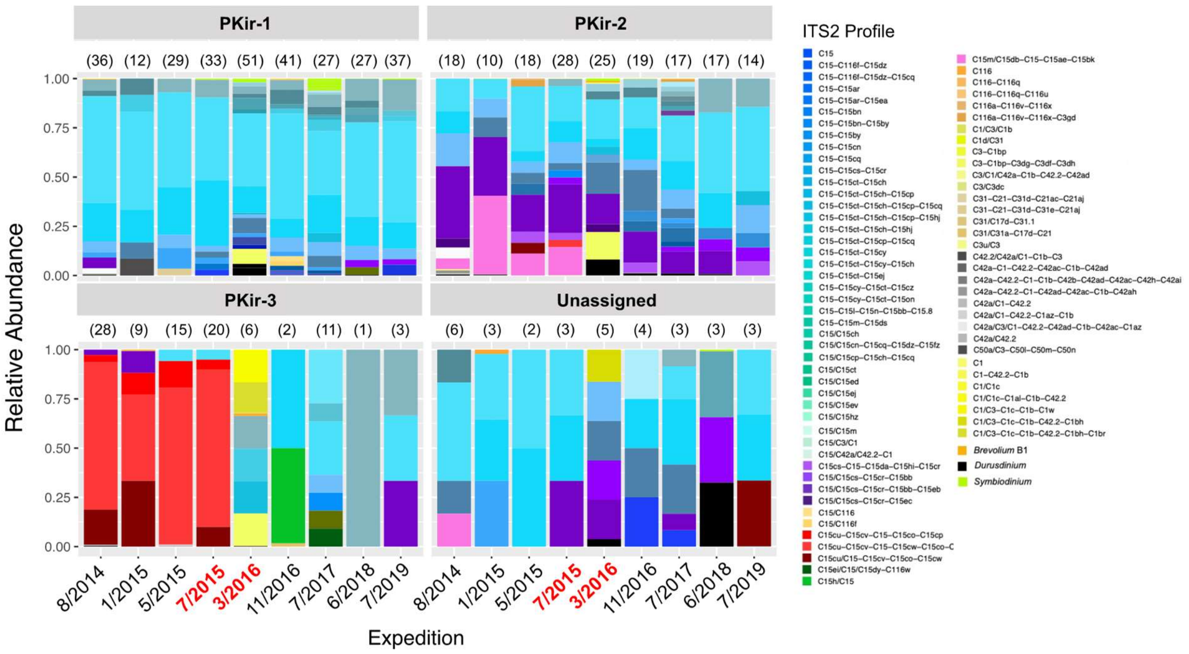


**Fig S7. Symbiodineaceae assemblages associated with each *Porites* lineage during each expedition.** Numbers above each bar indicate sample size.

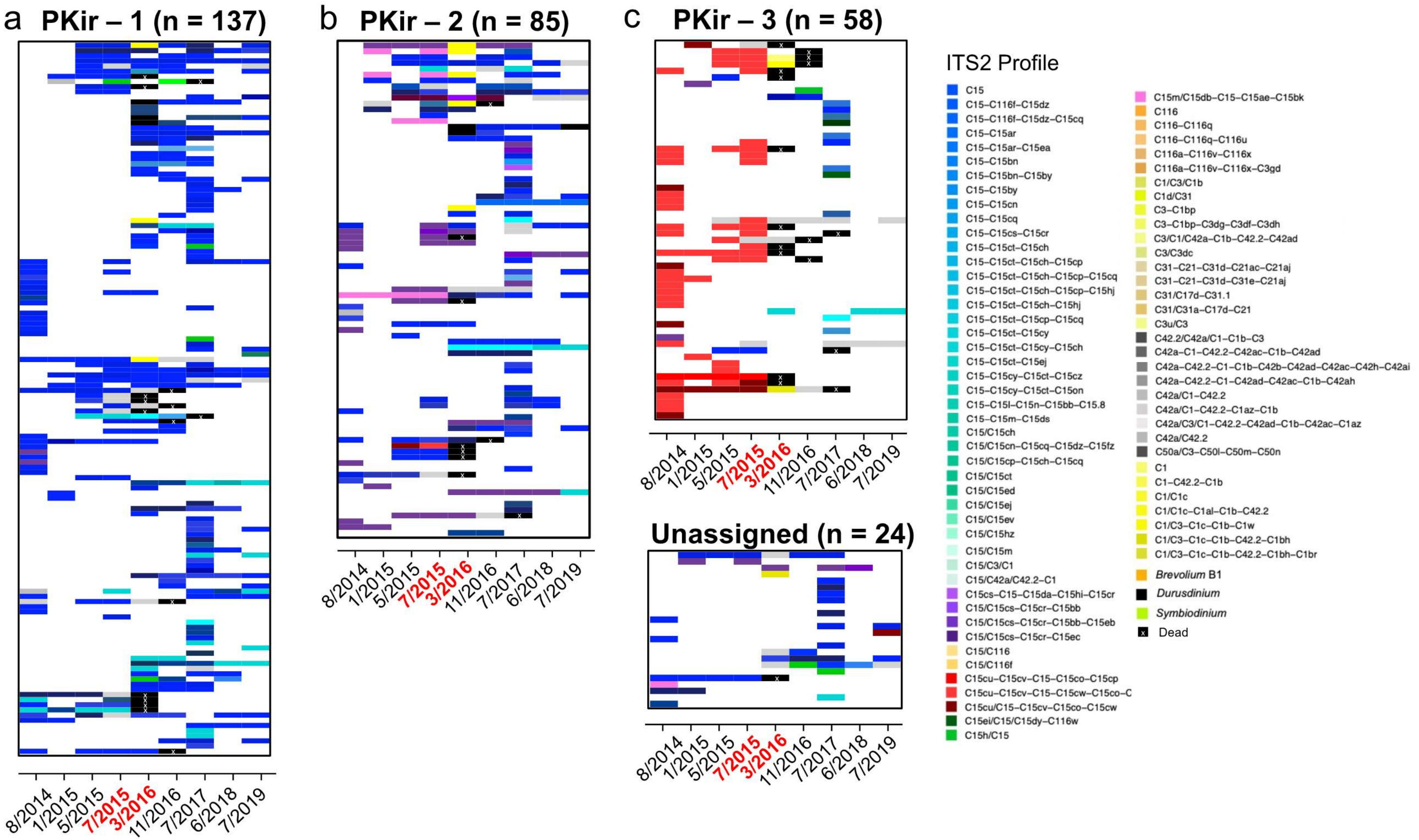


**Fig S8. Time series of Symbiodiniaceae associated with tracked colonies of each cryptic *Porites* lineage.** Each row represents an individual colony, with colour at each time point indicating the dominant symbiont profile. Colonies within each lineage are arranged in order of human disturbance (highest on top, lowest on bottom). Note that colonies that survived to 2017 were considered alive for survivorship analyses. All samples shown with >200 sequence reads (despite cut-off of >500 in main text analyses). Expeditions during the heatwave are shown in red text.

**Supp. File 1. Results of outlier analysis using colonies samples with 2bRAD**. See README sheet for more information. [Excel spreadsheet]

**Supp File 2. Additional information on genomic analyses.** See REAME sheet for more information [Excel spreadsheet]
